# Engineering chromatin to encode transcriptional immune memory in *Arabidopsis*

**DOI:** 10.64898/2026.09.10.750729

**Authors:** Linhao Xu, Jenia Binenbaum, Barbara Walkowiak, Harry Taylor, Vanda Adamkova, C. Jake Harris

## Abstract

Transcriptional memory enables organisms to respond more rapidly to recurrent stress, yet the underlying features of chromatin that contribute to this transcriptional recalibration remain poorly defined. Here we identify the genes displaying transcriptional memory in response to the bacterial immune elicitor, flg22, in Arabidopsis thaliana. In comparison to non-memory response genes, these memory genes show a preference for tissue-specific over uniform spatial expression patterning. The chromatin architecture of these genes in the resting state displays depletion of H3K4me3, elevation H3K27me3 and a subset are marked by H3K27me3-H3K4me3 bivalency. The H3K4me3 demethylase, JMJ14, is required for transcriptional memory, with JMJ14 occupancy enriched over memory gene loci. Upon priming, chromatin is reconfigured, with H3K4me3 levels increasing in a sustained manner at memory gene loci. To assess the function of this H3K4me3 accrual, we employ epigenome-engineering, observing that its targeted deposition at memory gene loci, including the WRKY29 locus, is sufficient to drive transcriptional memory and can endow plants with enhanced resistance to the bacterial pathogen, Pseudomonas syringae. Together, the findings demonstrate a causal role for H3K4me3 in transcriptional memory, under the regulation of JMJ14, and open the door for rational rewriting of chromatin to enhance organismal resilience.

## INTRODUCTION

While stimulus-induced responses are often considered reflexive and invariant, their magnitude and kinetics can be tuned by prior exposure, frequency, and environmental context^1^. Priming is one such example, whereby organisms can respond more rapidly and robustly to certain stresses if they have been previously challenged. This capability is widespread across eukaryotes^2,3^ and is associated with a range of physiological and metabolic processes, yet the underlying molecular basis is not well understood. In plants, priming is particularly prominent and can persist for days or significantly longer, with effects that can even be passed to subsequent generations^4^. These remarkable intergenerational effects are associated with global perturbation of the DNA methylation landscape^5^, although making direct links with functional genes can be difficult to establish^6,7^. Histone modifications are intimately associated with the transcriptional potential of genes *in cis*^8^, making chromatin an attractive substrate through which transcriptional events might be recorded on shorter, within generation timescales^9–12^.

Indeed, chromatin directly modulates transcription factor occupancy and polymerase engagement, thereby shaping stress-responsive transcriptional programs^13,14^. Chromatin states are not static, existing in dynamic equilibrium governed in part by the relative activities of histone modification writer and eraser machinery at any given locus^15,16^. Consistent with a role for chromatin in memory, early work showed that defense-associated WRKY transcription factor loci^17^ can acquire activation-associated histone modifications during priming, accompanied by enhanced inducibility upon subsequent challenge^18^. Subsequent studies have associated priming in both biotic and abiotic contexts with diverse chromatin features, including sustained H3K4 methylation^19–22^, altered regulation of Polycomb-associated H3K27me3^23–27^, and RNA polymerase II pausing^19,21^. Yet whether these features represent a passive consequence of previous transcriptional engagement, or affect future transcriptional responsiveness, remains largely unresolved. Furthermore, what distinguished genes capable of transcriptional memory from other response genes, and whether primed transcriptional response programs are deployed uniformly across plant tissues is yet to be resolved.

In plants, the perception the bacterial flagellin derived peptide, flg22, activates pattern-triggered immunity (PTI)^28^ and can confer enhanced resistance to subsequent infection^29^. Here we investigate the chromatin basis of priming during flg22-mediated PTI in *Arabidopsis thaliana*. We identify a suite of immune memory genes that show distinct chromatin features and display striking spatial specificity as compared to non-memory responsive genes. We identify chromatin regulators, including JMJ14 and CLF, required for this transcriptional memory, and upon priming, these memory genes acquire a sustained increase in H3K4me3. Using locus-specific epigenome engineering we show that deposition of H3K4me3 directly at endogenous genes is sufficient to enhance transcriptional responsiveness and increase pathogen resistance, demonstrating that memory-associated chromatin states can be rationally written.

## RESULTS

### Flg22 treatment establishes a transcriptional memory program

Flg22 is a component of bacterial flagellin that acts as a potent elicitor of pattern triggered immunity (PTI), initiating a well-characterised transcriptional program^30,31^. Flg22 is also a *bona fide* priming agent, as pre-treatment with flg22 confers enhanced resistance to bacterial pathogens^29,32^ (Supplementary Fig. 1). To investigate transcriptional memory in the context of plant immunity, we used flg22 exposure to induce transcriptional reprogramming in *A. thaliana*. To identify flg22 induced memory genes, we performed time-course RNA-seq (Fig. 1a) whereby five-day old seedlings were grown in liquid culture and treated with 25nM flg22 peptide (primed) or mock solution (naïve) for 2 hours, were allowed to recover for 3 days, and were then challenged with 100nM flg22. Whole seedlings were harvested at 5 minutes, 30 minutes, 3 hours, 24 hours and 48 hours after the second treatment for RNA-seq (Supplementary Fig. 2a). Differentially expressed genes showed strong overlap and tightly correlated expression dynamics with those reported by Bjornson et al., 2021^33^ (Fig. 1b and Supplementary Fig. 2c) confirming that our system recapitulates canonical flg22 responses. Principal component analysis (PCA) revealed clustering primarily by timepoint, with the 30-minute samples forming the most distinct group, and notably, the naïve and primed samples were clearly separated at this timepoint (Fig. 1c and Supplementary Fig. 2a).

**Fig. 1.**
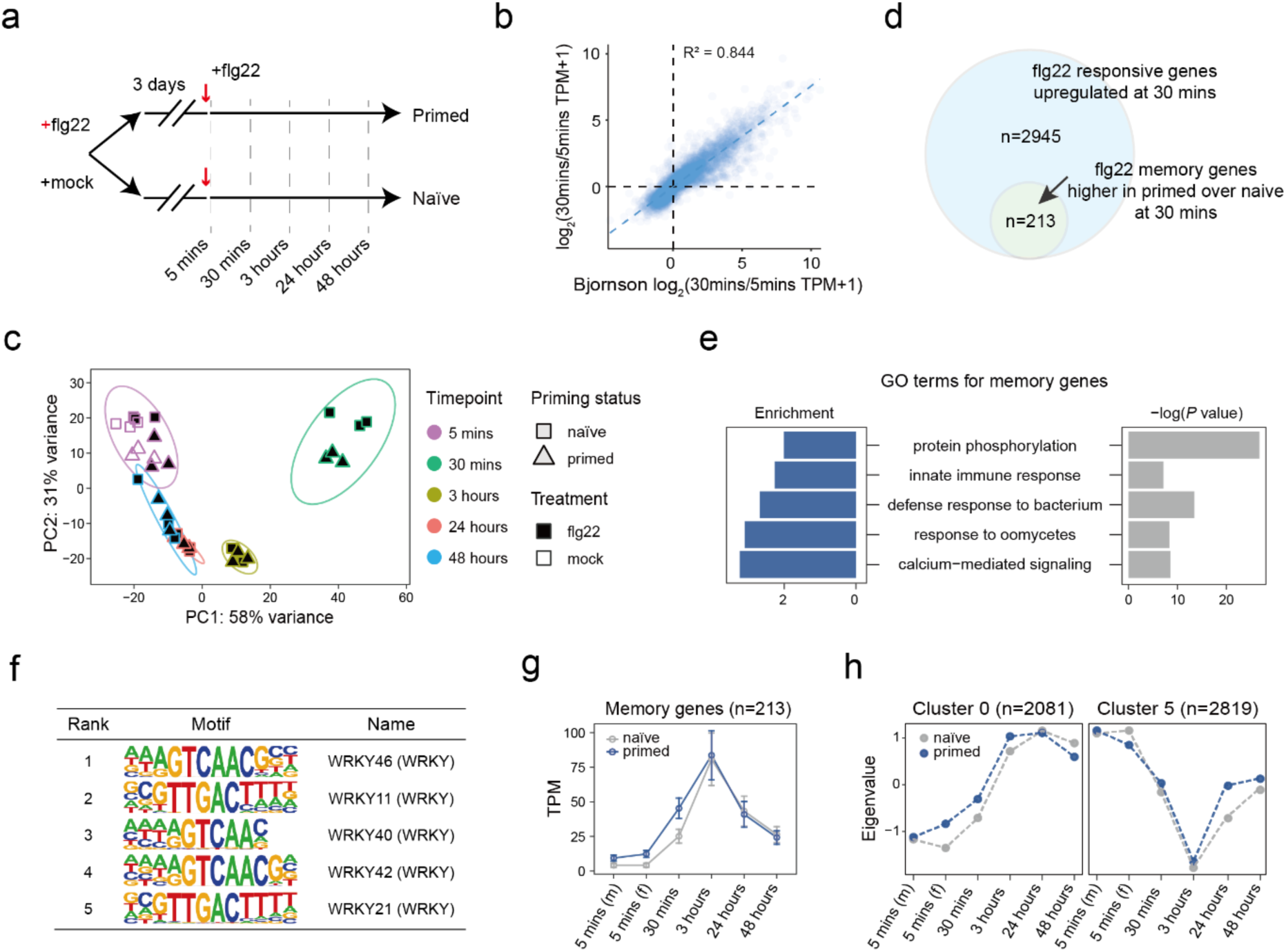
Flg22 induces a transcriptional memory gene program. **a.** Schematic of the experimental design for time-course RNA-seq. Seedlings were primed with 25 nM flg22 or mock treated, allowed to recover for 3 days, and challenged with 100 nM flg22. Samples were collected at 5 minutes (mins), 30 mins, 3 hours, 24 hours, 48 hours after the second treatment (n = 3 biological replicates per condition). **b.** Scatter plot showing the correlation between RNA-seq data generated in this study and the published dataset from Bjornson *et al*. (2021). **c.** PCA plot of transcriptomes collected at five time points after the recurrent stress (flg22) and recurrent mock treatment at 5 mins for both naïve and primed samples. Three biological replicates were sequenced for each time point. **d.** Venn diagram showing the overlap between flg22 responsive genes (n = 2945) and flg22 memory genes (n = 213). **e.** Gene Ontology (GO) enrichment analysis showing biological processes enriched among transcriptional memory genes. **f.** Motif enrichment analysis of transcriptional memory genes showing the top five enriched motifs. **g.** Line plots showing mean TPM expression levels (± SEM) of transcriptional memory genes under naïve and primed conditions across the indicated time points. **h.** Unbiased co-expression clustering of Cluster 0 (n = 2081) and Cluster 5 (n = 2819), showing expression dynamics under naïve and primed conditions across the indicated time points.

To increase discriminatory power at this key 30-minute timepoint, we repeated the experiment using 6 biological replicates per condition. We defined “memory genes” as those that were significantly more highly expressed in primed as compared to naïve samples at 30 minutes, and that were upregulated in response to flg22 at 30 minutes (Fig. 1d). This analysis identified 213 memory genes and showed strong overlap with the set detected in the original time course (Supplementary Fig. 2b), indicating that the memory-gene cohort is robust across experiments.

Memory genes were enriched for gene ontology (GO) terms related to biotic stress (Fig. 1e and Supplementary Fig. 3), and for encoding WRKY transcription factor binding motifs in their promoters (Fig. 1f) consistent with roles in pathogen defence^34,35^. Although memory genes showed higher induction in primed versus naïve at 30 minutes, their expression typically further peaked at 3 hours, reaching a similar level in both conditions (Fig. 1g). Compared to non-memory flg22-induced genes from the Bjornson dataset^33^, memory genes exhibited lower levels of basal expression and more delayed induction (Supplementary Fig. 2d). Priming therefore accelerates activation of weakly expressed responsive genes. Unbiased co-expression clustering revealed that the two largest gene cluster cohorts (C0, n = 2081 and C5, n = 2819) displayed more rapid induction, repression, and recovery in primed samples than in naïve (Fig. 1h and Supplementary Fig. 4), indicating that the stringently identified memory genes typify a much larger transcriptional program that follows a “primed response” trajectory.

### Single-nucleus transcriptomics reveals cell-type-specific activation of flg22 immune memory genes in the *Arabidopsis* shoot

Having identified memory genes based on their kinetic trajectories in whole seedlings, we wanted to establish where memory genes were expressed in the plant. While all tissues are thought capable of responding to flg22^29^, whether certain transcriptional programs are deployed in spatially distinct domains is largely unknown. We performed single nucleus RNA sequencing (snRNA-seq) on seedlings 30 minutes after flg22 treatment. After data filtering and quality control, the integrated dataset included 30,846 single-nuclei transcriptomes (Supplementary Fig. 5). Dimensionality reduction followed by UMAP projection showed that mock and flg22-treated nuclei were co-embedded, indicating that cell-type dominates the transcriptional identity space (Fig. 2a and Supplementary Fig. 6a).

**Fig. 2.**
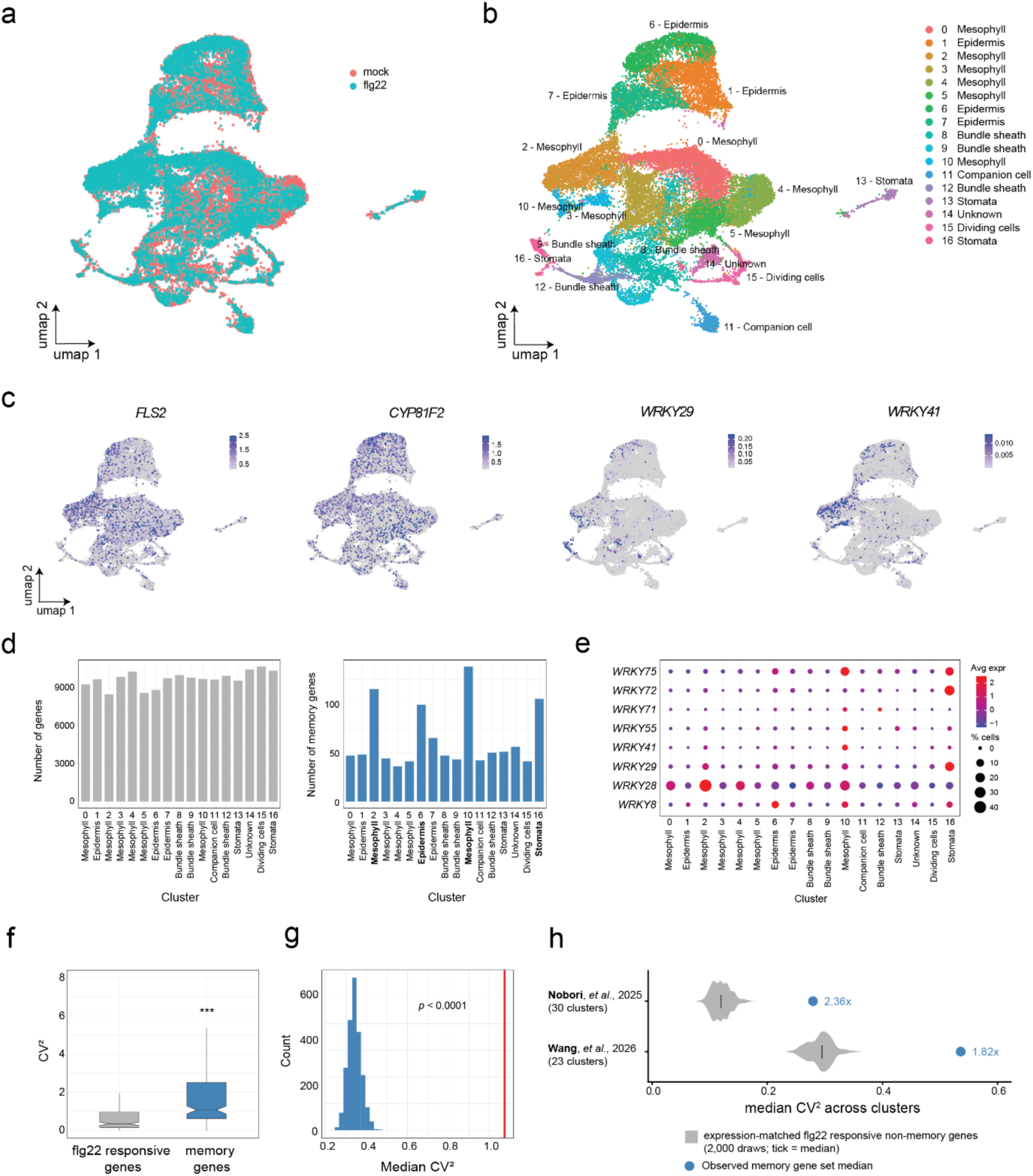
Flg22-induced activation of memory genes is restricted to a discrete subset of leaf cell types. **a.** UMAP visualization of single-nucleus RNA-seq data from Arabidopsis shoot tissue collected 30 mins after mock or flg22 treatment. Each point represents a single nucleus, colored by treatment (n = 2 biological replicates per condition). **b.** UMAP visualization of single-nucleus RNA-seq data colored by cell-type identity, revealing major shoot cell populations, including mesophyll, epidermis, bundle sheath, stomata, companion cells, and dividing cells. Cluster numbers are indicated. **c.** UMAP feature plots showing cell-type-resolved expression of the immune-associated genes *FLS2* and *CYP81F2* and the immune memory genes *WRKY29* and *WRKY41* across Arabidopsis shoot nuclei. **d.** Bar plots showing the distribution of all expressed genes (left) and memory genes (right) across single-nucleus cell clusters. The number of genes assigned to each cluster is indicated. **e.** Dot plot showing the expression patterns of *WRKY* immune memory genes across single-nucleus cell clusters. Dot color represents average normalized expression, and dot size indicates the percentage of nuclei expressing each gene within each cluster. **f.** Box plot showing the expression variability (coefficient of variation squared, CV²) between flg22-responsive genes and immune memory genes across single-nucleus clusters. Box plots show the median and interquartile range, with whiskers extending to 1.5× the interquartile range. Statistical significance was assessed using a Wilcoxon rank-sum test (*P* < 0.001). **g.** Permutation-based null distribution of the cross-cluster expression variability (CV²) calculated from randomly sampled sets of 213 flg22-responsive genes. The red vertical line indicates the observed median CV² of immune memory genes (1.07). Statistical significance was assessed using a Wilcoxon rank-sum test (*P* < 0.0001). **h.** Cross-cluster expression variability of transcriptional memory genes in independent single-nucleus (Nobori *et al.* 2025) and single-cell (Wang *et al*. 2026) transcriptomic datasets. Grey violin plots represent expression-matched null distributions generated from 2,000 random gene sets, with vertical ticks indicating the median. Blue points show the observed median CV² of memory genes, and fold enrichment over the null median is indicated.

Seventeen distinct transcriptional clusters were resolved (Fig. 2b and Supplementary Fig. 6b) and we performed automated annotation guided by known markers^36^ (see methods) revealing clusters belonging to all the major leaf cell types; mesophyll, epidermis, bundle sheath, companion, dividing and stomata (Supplementary Fig. 7). Comparing the expression of our memory genes to that of non-memory flg22-responsive genes, we observed that memory genes were often expressed in restricted clusters, whereas non-memory genes were more uniformly activated across clusters (Fig. 2c). Memory genes were preferentially expressed in clusters 2, 6, 10 and 16, corresponding to mesophyll, epidermal and stomatal subtypes (Fig. 2d). Given the enrichment of WRKY motifs in the memory gene set and their known role in plant defense^18^, we examined the *WRKY* genes that were themselves classified as memory genes in our RNA-seq dataset. Clusters 10 and 16 exhibited the highest proportion of *WRKY*-expressing nuclei together with the strongest average expression levels (Fig. 2e), reflecting the tissue-specific enrichment observed for memory genes more broadly.

To quantify the level of tissue specificity, we compared the expression variability of memory genes and non-memory flg22-responsive genes across clusters using the coefficient of variation squared (CV²). Memory genes exhibited significantly higher CV² values than the flg22 induced non-memory gene set (Fig. 2f). Performing a permutation test by repeatedly sampling 213 flg22-responsive genes (2,000 iterations) revealed that none of the random sets approached the CV² distribution of the memory genes (Fig. 2g). This suggests that memory genes are more spatially restricted than general flg22-responsive genes. To check whether this pattern was also observed in other datasets, we re-analyzed single-cell data from the recent Nobori *et al.,* 2025^37^ and Wang *et al.,* 2026^38^ studies. Excitingly, these revealed the same enhanced tissue specificity for the memory genes, as compared to non-memory flg22-responsive genes, which remained robust when controlling for expression-level distribution (Fig. 2h). Together, these results indicate that the transcriptional response of early immune memory is not evenly distributed across the shoot, and raises the possibility that certain cell types may hold designated memory functions.

### The chromatin signature of memory genes in the resting state features depletion of H3K4me3

To explore the chromatin landscape at memory genes, we integrated publicly available ChIP-seq and ATAC-seq datasets (see Sources of published data) and trained a boosted decision-tree model (XGBoost)^39^ using flg22 responsive non-memory genes as the background. This revealed that memory genes are relatively depleted for H3K4me3 and H3K36me3, and enriched for H2A.Z, chromatin openness (ATAC-seq signal) and H3K27me3 (Fig. 3a and Supplementary Fig. 8d, 9). Similar results were obtained when using transcriptionally equivalent non-memory genes as a background control (Supplementary Fig. 8a-c, 9). Plotting metaplot profiles of H3K4me3 and H3K27me3 -well-studied marks known to be positively and negatively, respectively, associated with transcriptional potential - also confirmed H3K4me3 depletion and slight 5’ enrichment of H3K27me3 (Fig. 3b). Together, these data indicate that memory genes possess a distinct chromatin profile, consistent with their low resting-state basal expression (Supplementary Fig. 2d), that includes depletion of euchromatic features such as H3K4me3 alongside enrichment of repression-associated H3K27me3.

**Fig. 3.**
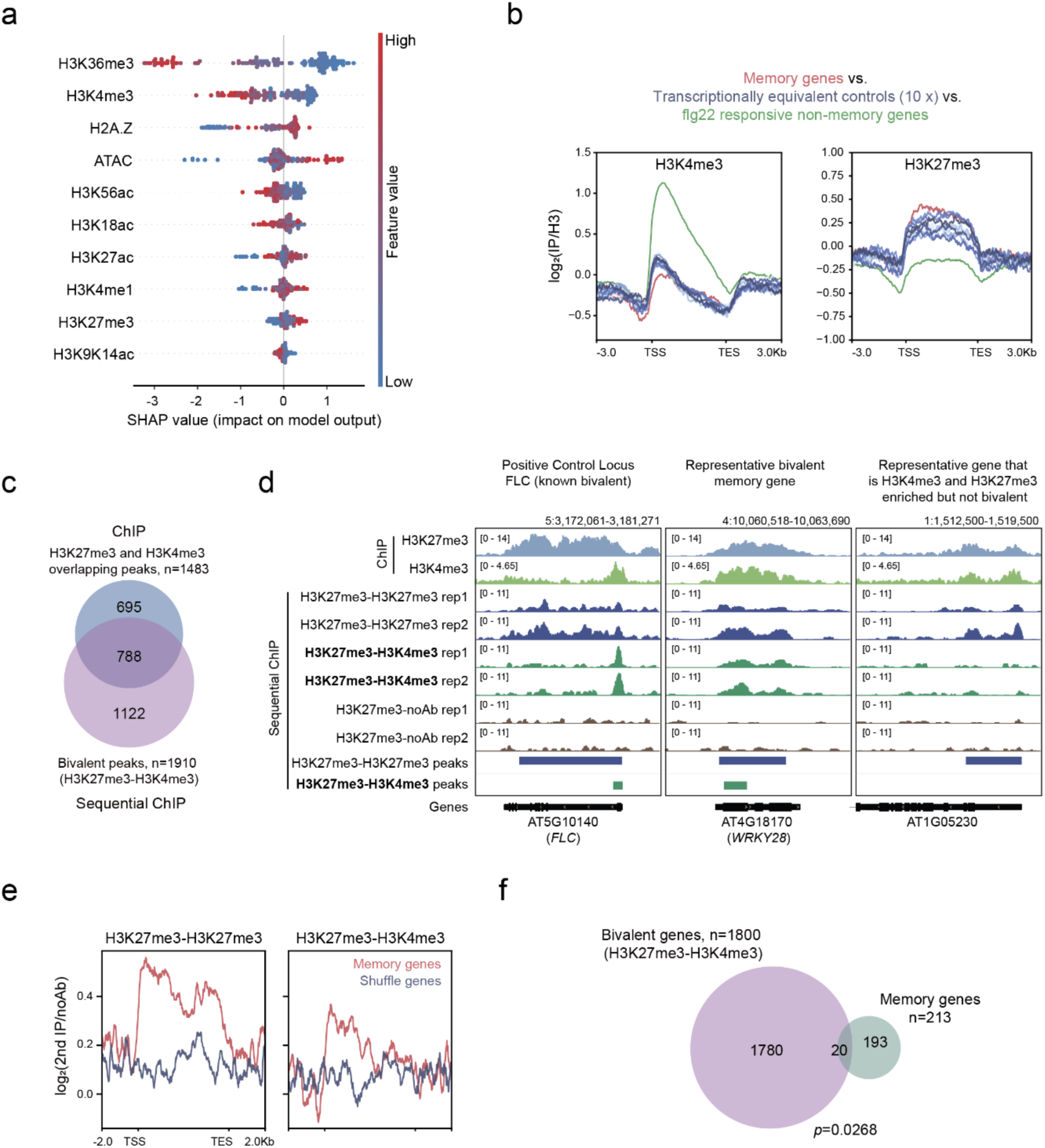
Chromatin features of transcriptional memory genes in the resting state. **a.** SHAP summary plot showing the contribution of chromatin features to an XGBoost model trained to distinguish memory genes from flg22-responsive non-memory genes. Features are ranked by importance, with SHAP values indicating their impact on model output and point color representing the relative feature value. **b.** Metaplots showing H3-normalized H3K4me3 and H3K27me3 profiles across memory genes (red), ten sets of transcriptionally equivalent non-memory control genes (blue), and flg22-responsive non-memory genes (green). **c.** Venn diagram showing the overlap between H3K4me3 and H3K27me3 overlapping peaks identified by conventional ChIP-seq (n = 1483) and bivalent H3K27me3-H3K4me3 peaks identified by sequential ChIP-seq (n = 1910). **d.** IGV genome browser snapshots showing H3K27me3, H3K4me3 and sequential ChIP-seq profiles at a positive-control bivalent locus (*FLC*), a representative bivalent memory gene (*WRKY28*), and a non-bivalent control gene (AT1G05230). **e.** Metaplots showing sequential ChIP-seq enrichment across memory genes (red) and shuffled control genes (blue) for H3K27me3-H3K27me3 (left) and H3K27me3-H3K4me3 (right). **f.** Venn diagram showing the overlap between H3K27me3-H3K4me3 bivalent genes (n = 1800) and flg22 memory genes (n = 213). Significance was assessed using a hypergeometric test.

Bivalent nucleosomes hold both activation and repression-associated chromatin marks, typically H3K4me3 and H3K27me3^40–42^, and are associated with genes poised for activation, particularly in developmental contexts^41,43^. We performed sequential ChIP to identify whether dual opposing marks co-occur on chromatin fragments at memory gene loci (Supplementary Fig. 10). This identified 1910 H3K27me3-H3K4me3 peaks, overlapping with >50% (788/1483) of peaks identified as being enriched for both marks individually using traditional ChIP-seq (Fig. 3c). Previously characterised bivalent loci, such as the floral repressor *FLC*^43^, were strongly H3K27me3-H3K4me3 enriched in our data, and genes overlapping with H3K27me3-H3K4me3 regions were highly GO-term enriched for floral organ and whorl development (Fig. 3d and Supplementary Fig. 10d), consistent with the previously described role for bivalency in development^44^. Together, these results demonstrate that the sequential ChIP-seq data provide a robust H3K27me3-H3K4me3 dataset to identify bivalent chromatin regions in *Arabidopsis thaliana*.

Over memory genes, there was a modest enrichment for H3K27me3-H3K4me3 signal with approximately only 10% of memory genes directly overlapping with bivalent peak regions (Fig 3e,f). While this overlap is higher than expected by chance (*p* = 0.0268), it indicates that only a subset of memory genes holds canonical bivalency.

### JMJ14 and CLF are required for transcriptional memory

Next, we employed a reverse genetics approach to identify chromatin pathways that might play a role in transcriptional memory. A range of nucleosome modification mutants (including mutants defective in Polycomb, H2A.Z deposition, H3K4me3 and H3K27me3, see Supplementary Fig. 11) were assessed for their impact on flg22-induced transcriptional memory. While the majority of mutants had little impact on the transcriptional dynamics of the memory genes selected, the H3K27me3 methyltransferase mutant, *clf*, displayed a highly reduced transcriptional memory phenotype (Supplementary Fig. 11b). The defect was confirmed by full transcriptome analysis, whereby memory gene expression was impaired after the second stimulus in *clf* as compared to Col-0 (Supplementary Fig. 11c). *clf* mutants are known to have pleiotropic effects on development^45^ and while these phenotypes were not visible at the early developmental timepoint used in these assays (Supplementary Fig. 12), we cannot exclude the possibility of indirect effects.

Another mutant that appeared perturbed in transcriptional memory at representative loci was *jmj14* (Supplementary Fig. 11b). JMJ14 encodes an H3K4me3 demethylase protein^46^. Transcriptome analysis of *jmj14* confirmed a near complete loss of transcriptional memory as compared to WT (Fig. 4a,b). We compared levels of H3K4me3 in *jmj14* to WT^47^, which confirmed that JMJ14 prunes H3K4me3 primarily at the 5’ end of gene bodies where H3K4me3 is most enriched (Fig. 4c,d and Supplementary Fig. 13). The occupancy of JMJ14 itself was highly enriched over memory genes as compared to transcriptionally equivalent control gene sets (Fig. 4c,d and Supplementary Fig. 13), indicating that JMJ14 specifically targets memory gene loci. Together these data indicate that both polycomb and H3K4me3 pathways play an important role in transcriptional memory, with JMJ14 targeting memory genes for removal of H3K4me3, consistent with the observed depletion of H3K4me3 in the resting state (Fig. 3a,b).

**Fig. 4.**
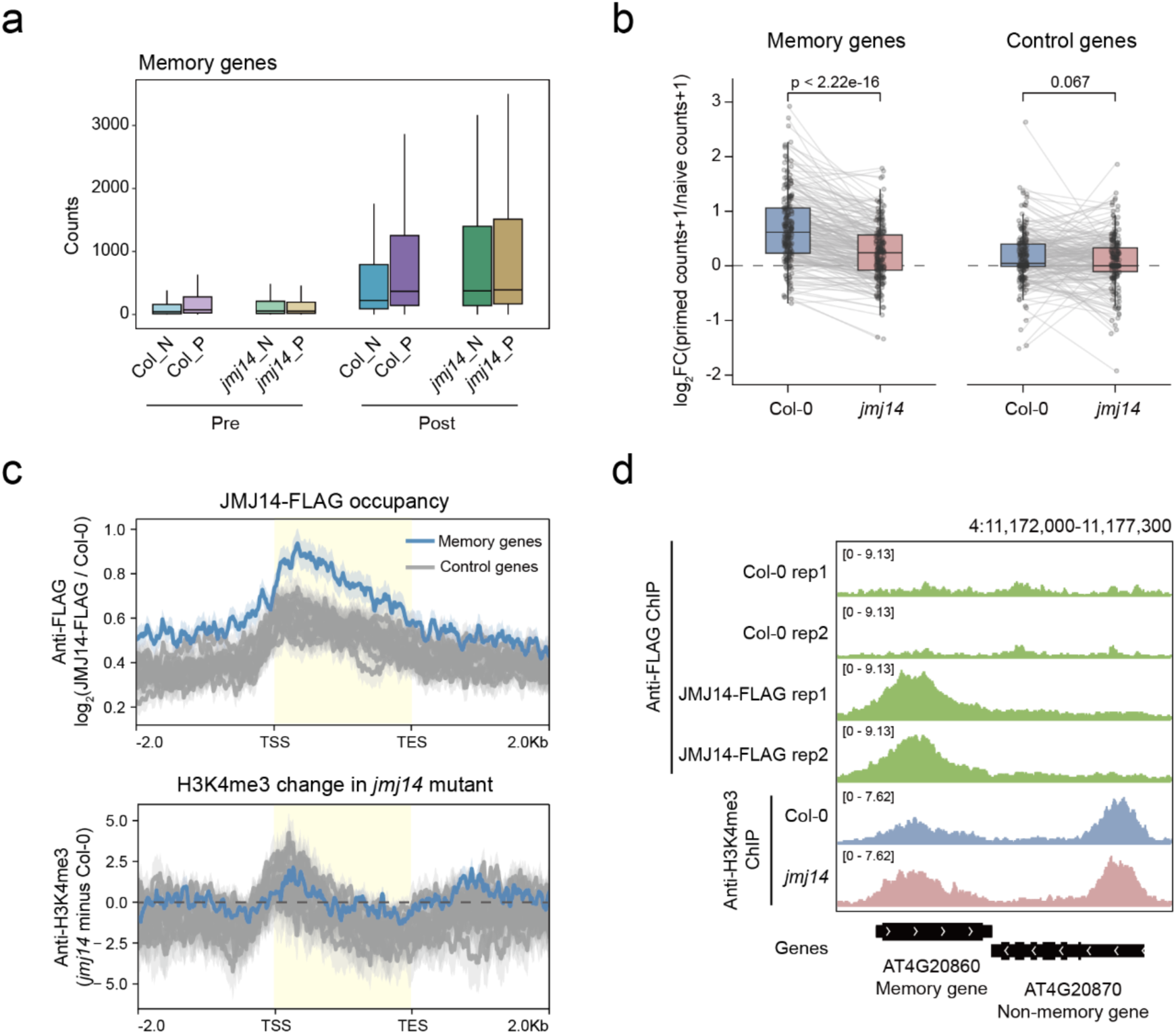
JMJ14 is required for flg22-induced transcriptional memory. **a.** Box plots showing RNA-seq read counts for memory genes in Col-0 and *jmj14* under naïve (N) and primed (P) conditions before (Pre) and following recurrent flg22 challenge (Post). Pre samples contained three biological replicates per condition, and Post samples contained six biological replicates per condition. Box plots show the median and interquartile range, with whiskers extending to 1.5x the interquartile range. **b.** Paired box plots showing the priming response, calculated as log_2_[(primed counts + 1)/(naïve counts + 1)], for memory genes and control genes in Col-0 and *jmj14*. Grey lines connect the same genes between genotypes. Box plots show the median and interquartile range, with whiskers extending to 1.5x the interquartile range. *P* values were calculated using a paired Wilcoxon signed-rank test. **c.** Metaplots showing JMJ14-FLAG occupancy (top) and changes in H3K4me3 in the *jmj14* mutant (bottom) across memory genes (blue) and ten sets of transcriptionally equivalent non-memory control genes (grey). ChIP-seq data from (GSE204681)^47^. **d.** IGV genome browser snapshots showing JMJ14-FLAG occupancy and H3K4me3 profiles at a representative memory gene (AT4G20860) and a neighboring non-memory gene (AT4G20870).

### Priming induced chromatin reconfiguration features gain of H3K4me3

Next, we asked how the chromatin state is altered upon priming. At the end of the 3-day recovery period (as outlined in Fig. 1a), when flg22-induced transcription has largely recovered (Fig. 1g), we collected samples for chromatin profiling, performing ATAC-seq (chromatin accessibility) and ChIP-seq for nine histone modifications and Pol II (Ser5P form and total Pol II) in both naïve and primed plants (Supplementary Fig. 14-16). While most marks showed relatively limited memory-gene-specific effects, H3K4me3 stood out as significantly increased in the primed state (Fig. 5a and Supplementary Fig. 17).

**Fig. 5.**
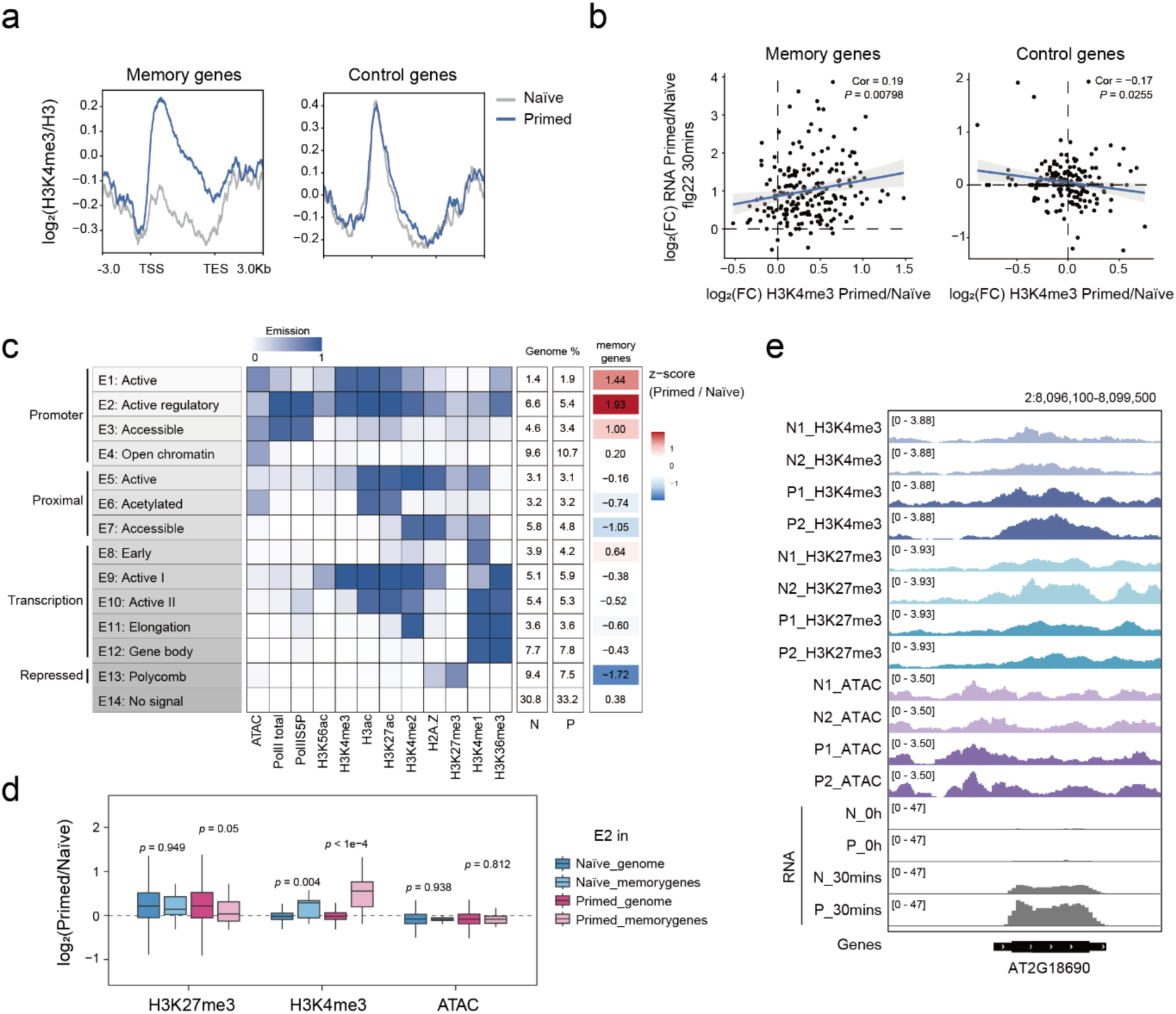
Priming-induced chromatin reconfiguration is marked by H3K4me3 gain. **a.** Metaplots showing H3-normalized H3K4me3 profiles across memory genes (left) and one set of transcriptionally equivalent non-memory control genes (right) under naïve (gr y) and primed (blue) conditions after 3 days of recovery and before recurrent flg22 treatment. Profiles were generated from ChIP-seq data merged from two biological replicates per condition. **b.** Scatter plots showing the relationship between priming-induced changes in H3K4me3 and transcriptional output at 30 mins after recurrent flg22 treatment for memory genes (left) and one set of transcriptionally equivalent non-memory control genes (right). H3K4me3 changes are shown as log_2_fold change (primed/naïve), and RNA changes as log_2_fold change (primed/naïve). Pearson correlation coefficients and *P* values are indicated. **c.** ChromHMM emission heatmap showing the chromatin features defining 14 chromatin states at the 3-day recovery stage before recurrent flg22 treatment, grouped into promoter, proximal, transcriptional, and repressed categories. Each row represents a chromatin state, while the columns show the emission probabilities of individual chromatin features, the proportion of the genome assigned to each state under naïve (N) and primed (P) conditions, and a z-score heatmap showing the primed/naïve change in state abundance. **d.** Boxplot showing log_2_ fold-change values (primed/naïve) of H3K27me3 and H3K4me3 ChIP-seq signals and ATAC-seq signals for E2 intervals across genome and memory genes under both naïve and primed conditions. The boxplot’s top, mid-line, and bottom represent the upper quartile, median, and lower quartile, respectively. *P* values were calculated using Wilcoxon rank-sum tests and are indicated above the comparisons. **e.** IGV genome browser snapshots showing H3K4me3, H3K27me3, and ATAC-seq profiles from individual biological replicates under naïve and primed conditions, together with RNA-seq signal (mean of six biological replicates) immediately before (0 h) and 30 mins after recurrent flg22 treatment, at the representative memory gene AT2G18690. N1, N2 are naïve replicates; P1, P2 are primed replicates.

As H3K4me3 is associated with active transcription and we observed increased H3K4me3 in primed plants, we next asked whether this elevation was associated with enhanced transcriptional potential. We examined the relationship between changes in H3K4me3 between naïve and primed, and the transcriptional output at 30 minutes. We observed a slight but significant positive correlation (Pearson r = 0.19, *p* = 0.008), which was not observed for control loci (Fig. 5b). This is consistent with previous reports showing that H3K4me3 persists after transcription has returned to baseline in the context of heat stress memory^20^, suggesting that it could help to facilitate transcriptional enhancement.

Next, we assessed how overall chromatin state reconfigures upon priming, beyond focusing on individual marks in isolation^48–50^. Using a Hidden Markov based modelling approach (ChromHMM^51,52^), our data uncovered a 14-state model that we selected for downstream analysis (Supplementary Fig. 18a). Chromatin states were annotated based on histone mark enrichments, genomic distributions, and associated expression levels (Fig. 5c and Supplementary Fig. 18b). In total, four major groups were defined: promoter-, proximal-, transcription-associated and repressed states. The promoter group contained the three chromatin states that showed the highest transition dynamics between naïve and primed over memory gene loci. Of these, E2 - a state characterized by modest accessibly, activation associated mark enrichment and modest levels of repression associated H3K4me2 and H3K27me3 -showed the highest level of state transition with priming (Fig. 5c). Further resolving the chromatin dynamics in the E2 state, we observed significant increases of H3K4me3 in the primed state, with H3K27me3 trending towards reduction at loci overlapping with memory genes (Fig. 5d,e and Supplementary Fig. 19). Together, these data underscore that the chromatin state underlying memory gene responsiveness involves dynamic gain of H3K4me3 as a primary feature, along with chromatin containing both activation and repression associated features.

### Targeted deposition of H3K4me3 at the *WRKY29* promoter primes gene expression and pathogen resistance

Having found that H3K4me3 accumulates at memory genes in the primed state, we were motivated to assess whether the mark is causal for transcriptional memory. *WRKY29* is a transcription factor that is known to play a key role in plant defense^35^, and was identified here as a memory gene. Previously, *WRKY29* was shown to display transcriptional memory and acquire euchromatic marks in response to pathogen priming^18^. However, whether these marks directly facilitate future transcriptional potential is unknown. To address this, we employed our recently described CRISPR/dCas9-based system for locus-specific H3K4me3 deposition, SunTag:SDG2^53^, to target *WRKY29*. We designed 3 guides targeting the promoter region of *WRKY29*. ChIP-seq revealed highly specific enrichment of the SunTag at this locus (Fig. 6a and Supplementary Fig. 20), accompanied by increased H3K4me2/3 in the surrounding region (Fig. 6a,b and Supplementary Fig. 21a).

**Fig. 6.**
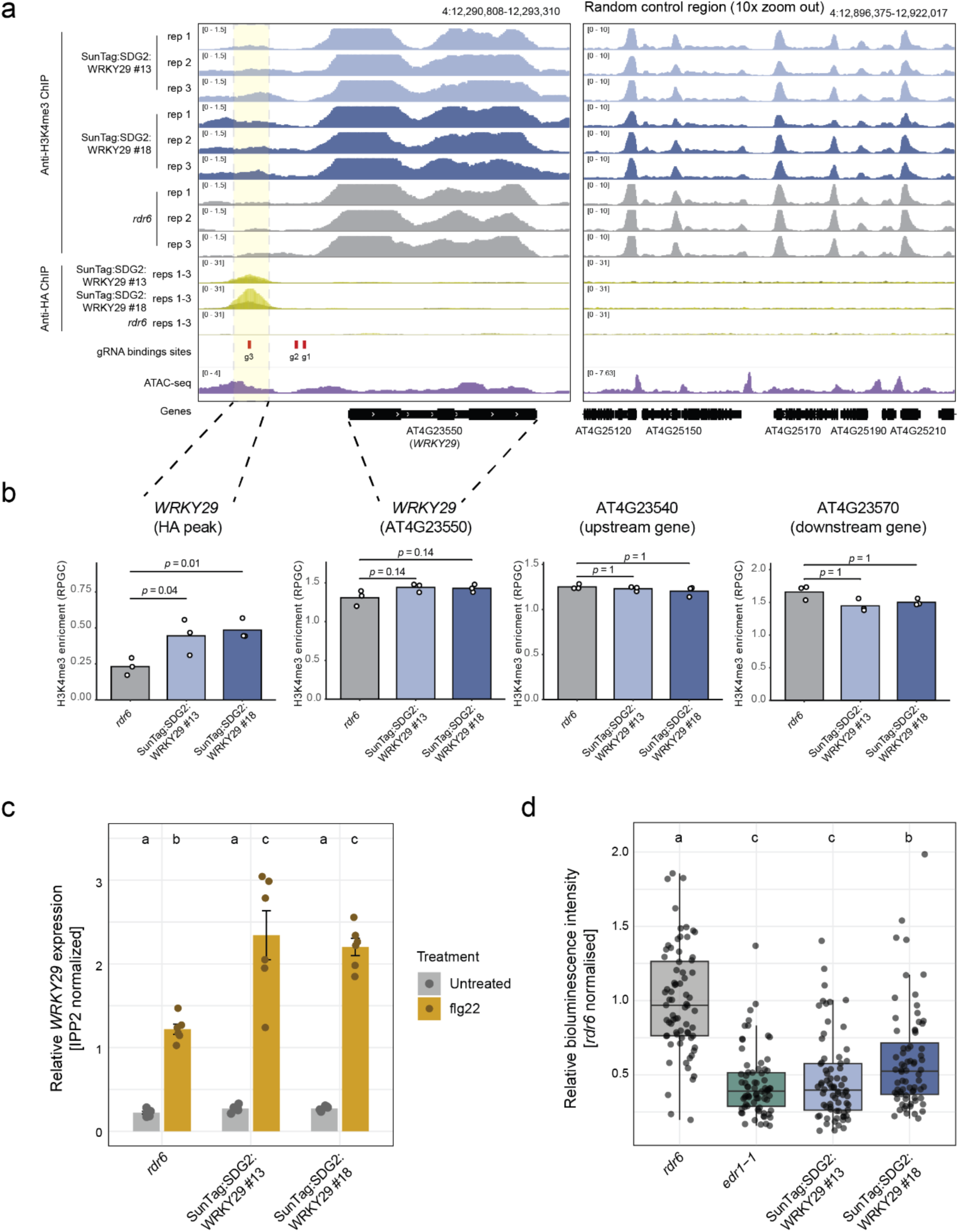
Targeted H3K4me3 deposition at WRKY29 is sufficient to synthetically prime transcriptional memory and enhance pathogen defense. **a.** IGV genome browser snapshots showing H3K4me3 enrichment at the *WRKY29* locus in two independent SunTag:SDG2:WRKY29 lines (#13 and #18) and the *rdr6* control, together with anti-HA ChIP-seq signal marking SunTag occupancy, gRNA-binding sites, and ATAC-seq accessibility. Three biological replicates are shown for H3K4me3 ChIP-seq, while the anti-HA ChIP-seq signal represents the overlay of three biological replicates. The yellow shaded region indicates the targeted gRNA-binding region. A random control region is shown on the right. **b.** Bar plots showing H3K4me3 enrichment (RPGC), at the *WRKY29* HA-peak region, the *WRKY29* gene body (AT4G23550), and the upstream (AT4G23540) and downstream (AT4G23570) control genes in *rdr6* and two independent SunTag:SDG2:WRKY29 lines (#13 and #18). Individual biological replicates are shown as points. *P* values were calculated using one-sided Welch’s t-tests testing whether each line had greater enrichment than *rdr6*, with Holm-adjusted *P* values indicated above the comparisons. **c.** Bar plots showing IPP2-normalized *WRKY29* expression in *rdr6* and two independent SunTag:SDG2:WRKY29 lines (#13 and #18) under untreated and flg22-treated conditions (n = 6 biological replicates). Bars represent mean expression with individual biological replicates shown as points. Different letters indicate statistically significant differences between groups, determined by one-way ANOVA followed by Tukey’s HSD test (*P* < 0.05). d. *Pst*::LUX bioluminescence assay quantifying bacterial colonization at 3 days post-inoculation in *rdr6*, two independent SunTag:SDG2:WRKY29 lines (#13 and #18), and the constitutively primed control genotype *edr1-1*. Box plots show the median and interquartile range, with whiskers extending to 1.5x the interquartile range; individual wells are shown as points. Different letters indicate statistically significant differences between genotypes, determined by one-way ANOVA followed by Tukey’s HSD test (*P* < 0.05). n = 96 wells, with three seedlings per well.

Using these lines, we then assessed transcript kinetics at the *WRKY29* locus in response to flg22. In the untreated condition, the basal levels of *WRKY29* were virtually indistinguishable between the SunTag:SDG2 lines and the background (*rdr6*) controls (Fig. 6c). The *rdr6* background is used as it has previously been shown to support higher expression and reduced silencing of the SunTag constructs^53,54^, and we do not observe any impact on transcriptional memory in these lines (S. Fig. 11b). However, when treated with flg22 the SunTag:SDG2 lines showed a significantly elevated response at 30 minutes, with *WRKY29* expression levels approaching nearly double that of the non-transgenic controls (Fig. 6c), thereby recapitulating the transcriptional memory phenotype. Importantly, a control SunTag:SDG2 line targeting a different locus, *FWA*^53^ did not show the same effect (Supplementary Fig. 22a), indicating that site-specific deposition of H3K4me3 is required to induce transcriptional memory at *WRKY29*. We also assessed a catalytically dead control^53^, to determine whether deposition of H3K4me3 is required, or whether simply occupancy of SunTag:SDG2 system at the *WRKY29* promoter is responsible for the enhanced transcriptional response. This version also had no effect on transcriptional memory, despite confirmation of its occupancy at the same position in the promoter (Supplementary Fig. 22b,c). Together, the data strongly indicate that transcriptional memory at *WRKY29* can be directly mediated by H3K4me3.

As *WRKY29* is upregulated in response to the microbial elicitor, flg22, and priming is typically associated with enhanced pathogen resistance^29^, we challenged these *WRKY29* targeting lines with a bioluminescent strain of *Pseudomonas syringae* DC3000 (*Pst::LUX*)^55^, finding that they displayed enhanced resistance (Fig. 6d). To assess whether enhanced loading of H3K4me3 at *WRKY29* affects priming potential, we pre-treated the SunTag:WRKY29 lines with flg22, finding that they displayed further enhanced transcriptional memory effects (Supplementary Fig. 23). Finally, we targeted two additional memory gene loci, *WRKY41* and *RMG1*^56^, with SunTag:SDG2 finding that these genes also display transcriptional memory (Supplementary Fig. 24).

Therefore, our results demonstrate that deposition of H3K4me3 can generally be used to drive transcriptional memory and can endow plants with improved priming-associated pathogen performance.

## DISCUSSION

Here we investigate the chromatin basis of transcriptional memory under immune stimulation in *Arabidopsis thaliana*, using both loss-and gain-of-function approaches to test causal function. The work underscores a central role for H3K4me3 in the process, allowing us to propose a general model (Fig. 7). Briefly, memory genes undergo continuous H3K4me3 turnover in the resting state, with JMJ14-mediated removal maintaining low levels of H3K4me3. Priming transiently activates these genes, leading to co-transcriptional H3K4me3 acquisition that persists after stimulus withdrawal. This H3K4me3 accrual, which we can recapitulate through locus-specific epigenetic editing, is sufficient to drive the transition to a poised transcriptional state.

**Fig. 7.**
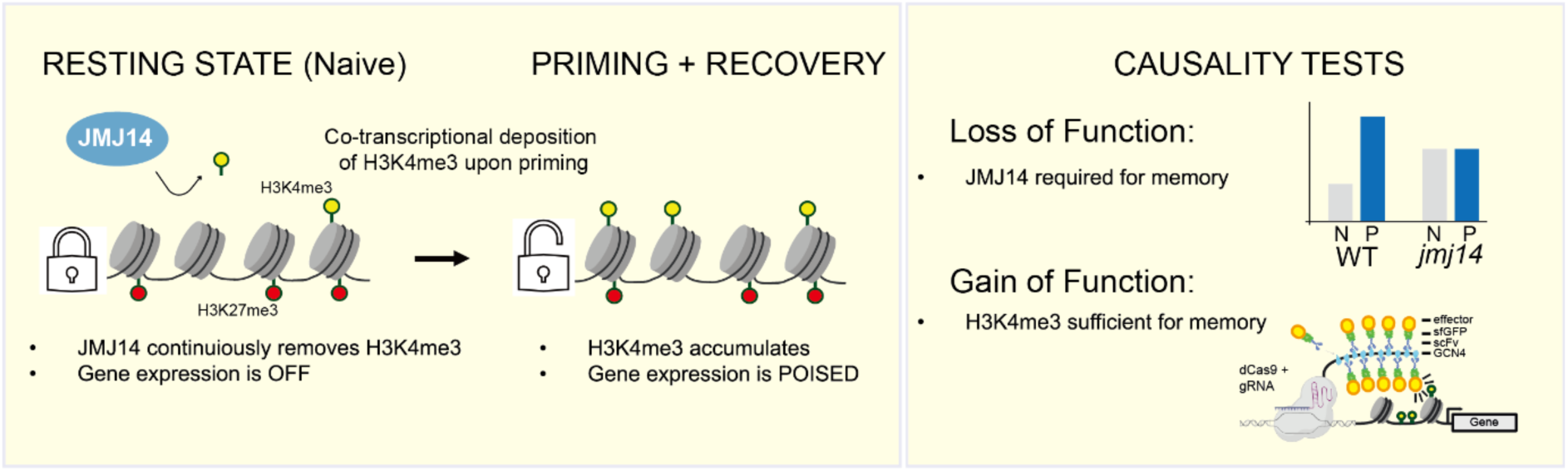
Proposed model for chromatin reconfiguration underlying flg22 transcriptional memory. JMJ14 maintains low H3K4me3 levels at memory genes in the resting state. Following priming, H3K4me3 accumulates and persists during recovery, contributing to enhanced transcriptional responsiveness upon recurrent flg22 treatment. Loss-and gain-of-function experiments support a causal role for H3K4me3 dynamics in transcriptional memory.

In *Arabidopsis thaliana*, there are 21 JMJ-domain containing histone demethylase proteins^57^. Our work revealed a critical role for H3K4me3 demethylation, mediated by JMJ14, in PTI priming (Fig 4). This is consistent with previous work showing that H3K4me3 demethylases play a key role in a range of stress responses, both biotic^58^ and abiotic^59^. Interestingly, JMJs targeting H3K27me3 have also been implicated in priming and stress responses^25,59^. For instance, recent work showed that REF6 is required to clear H3K27me3 at certain defense loci to maintain transcriptional responsiveness upon flg22 stimulation^60^. While we observed modest enrichment of H3K27me3 over memory genes (Fig. 3a,b), we did not observe appreciable loss of this mark in the primed state (Supplementary Fig. 17). This is consistent with recent independent studies on cold acclimation^61,62^ showing that loss of H3K27me3 is generally uncoupled from transcriptional stimulation, and early seminal work showing that H3K27me3 islands become fractionated rather than appreciably removed, in response to drought priming^63^. Given the well-established role of JMJs in resetting the memory of cold at *FLC*^64^, our results underscore the importance of histone methylation removal in establishing memory-competent chromatin states and in the future it will be important to understand how JMJ14 is regulated during flg22-priming.

There are many chromatin states that could afford poised transcriptional potential, including bivalency^44,65^. Our work here identified canonical H3K27me3-H3K4me3 bivalency occurring at a small subset (10%) of memory genes. Therefore, while canonical bivalency may be important at specific loci, it is unlikely to represent a general feature of immune memory genes. Individually, however, both H3K27me3 and H3K4me3 were found to play critical roles, with both the *clf* (H3K27me3 deposition) and *jmj14* (H3K4me3 removal) mutants displaying drastically reduced transcriptional memory potential (Fig. 4 and Supplementary Fig. 11). Non-canonical bivalency, H3K27me3-H3K18ac, was recently found to be associated with certain flg22-induced camalexin-response genes^66^, and so it remains possible that bivalency in this non-canonical form is a more widely-endowed feature of memory genes in plants.

Priming is often thought to be systemic across tissues in plants, mediated by non-cell autonomous effects such as hormone signalling^1,67^. Our work here highlights the potential of individual cells to encode memories of their life-history, and the single-nuclei transcriptomes indicate that memory is likely non-uniform across tissues. This cell-specificity suggests that more pronounced chromatin rearrangements may be masked by the bulk profiling approaches used here. It is also important to note that our single nucleus RNA-seq data was performed after acute treatment of flg22, so whether priming affects the nature and distribution of cells that respond upon subsequent flg22 exposure will be important to establish in future work, ideally alongside at chromatin profiling at this resolution. However, together with recent studies showing ‘gated’ transcriptional responses^38^ that are spatially restricted to cells adjacent to *Pseudomonas* colonisation sites^37^, the results suggest that immune signalling and memory are fundamentally heterogeneous, even within individual tissues^68–71^.

Many chromatin correlates of priming have been identified, yet direct tests of causality remain rare^12,72,73^, particularly outside of DNA methylation-based mechanisms^74^. Here, we show that targeted deposition of H3K4me at the *WRKY29* locus is sufficient to confer a priming-like response. As we also observe a concomitant increase of both H3K4me2/3 in these lines - consistent with SDG2’s biochemical capacity^75^ - we cannot exclude a possible contribution from H3K4me2. However, as H3K4me2 is more associated with negative transcriptional potential^76,77^, and is not observed to increase at this locus or at memory genes in general during priming (Supplementary Fig. 17, 21b), we favour that H3K4me3 is driving this effect. *WRKY29* is a particularly informative locus, having been originally identified as a pathogen memory gene that acquires activation-associated chromatin marks upon priming, while remaining transcriptionally silent until re-challenged^18^. Consistent with this, targeted H3K4me3 deposition enhances inducibility without driving constitutive expression (Fig. 6c) indicating that H3K4me3 can prime transcriptional potential^19^ rather than directly activate transcription as it does at other loci^53,78^. Importantly, we observe a similar effect at two other loci, *RMG1* and *WRKY41*, suggesting that the effect may be generalisable across memory genes. *RMG1* was previously identified as a DNA methylation-sensitive flg22-inducible gene^56^, raising the possibility that SDG2 targeting also primes this locus by antagonising DNA methylation establishment^78^.

Recent data from mammals suggests that while H3K4me3 can be dispensable for transcriptional maintenance^79,80^, it also supports transcriptional stimulation in permissive chromatin environments^81,82^. Our data highlights a key role for H3K4me3 in the direct establishment of a permissive state that facilitates future transcriptional output. Together with the evidence that H3K4me3 can be maintained through DNA replication^83–85^, these findings suggest a dual role for H3K4me3 in both encoding transcriptional history and enhancing transcriptional response. As the use of targeted epigenome engineering rapidly expands across both plant and animal research^54,81,86–92^, the tools are being deployed to answer questions across a wide of chromatin biology, from gene expression^74^ to recombination control^93–95^.

Our epigenome-engineered plants showed an additive effect with priming (Supplementary Fig. 23), indicating that although H3K4me3 is sufficient to increase transcriptional responsiveness, there are likely other features that contribute. Consistent with this, the chromatin state showing the largest priming-associated transition, E2, was defined by the co-occurrence of multiple chromatin modifications (Fig 5). Co-recruitment of distinct chromatin modifiers may therefore provide a potent means of reconfiguring target loci^96,97^. As natural priming can impose a fitness cost^98^, developing epigenome-editing tools that minimise off-target activity^99,100^ and afford precise spatial and temporal control will be essential for dissecting basic chromatin function and for improving crop performance.

## MATERIALS AND METHODS

### Plant materials and growth conditions

All *Arabidopsis thaliana* plant materials used in this work are Columbia (Col-0) ecotype background. The *clf-81* mutant was provided by Dr. Matthew Naish, the *atx1-4* (SALK_140755) and *jmj14* (SALK_135712C) mutants were provided by Dr. Régis Lopes Corrêa, the *arp6-1* (SAIL_599_G03) and *h2a.z-5*^101^ mutants were provided by Prof. Israel Ausin, the *edr1-1* mutant^55^ was provided by Prof. Juriaan Ton and the *rdr6-15* (SAIL_617_H07) was previously described^54^. T-DNA mutant *lhp1-6* (SALK_011762) was obtained from Nottingham Arabidopsis Stock Centre (NASC). Plants grown on plates were sown onto 1/2 x Murashige-Skoog (MS) medium, 1% sucrose, and 0.8% agar (pH 5.7), stratified for 2 days at 4 °C in the dark, then transferred to growth chambers (Percival CU-41L4D) at 21 °C under long day light conditions (16 h light/8 h dark). Plants grown on soil were grown in F2 soil at 20 °C under long day light conditions (16 h light/8 h dark).

### Plasmid construction

SunTag:SDG2:WRKY29_g1g2g3 was generated following a previously described protocol^102^. Briefly, the WRKY29_g1 RNA cassette was PCR-amplified and inserted into KpnI (NEB, R3142L)-digested SunTag:SDG2 (no-guide)^53^ using In-Fusion (Takara, 638948) cloning. The g2 and g3 RNA cassettes were added sequentially using the same strategy. The WRKY29 g1, g2 and g3 target sites were located 118, 172 and 489 bp upstream of the transcription start site (TSS), respectively. The catalytically dead SDG2 control (dSDG2) was described previously^53^. SunTag:SDG2:WRKY41_g1g2g3 and SunTag:SDG2:RMG1_g1g2g3 constructs were generated using the same cloning strategy. The WRKY41 g1, g2 and g3 target sites were located 36, 132 and 213 bp upstream of the TSS, respectively, whereas the RMG1 g1, g2 and g3 target sites were located 1062, 1938 and 2840 bp upstream of the TSS, respectively. All transgenic plants were generated by *Agrobacterium tumefaciens* strain AGL1 using floral dip^103^. Guide RNA sequences are listed in Supplementary Table 1.

### Priming treatment

Surface-sterilized seeds were sown into 1 x MS Murashige-Skoog (MS) liquid medium (1.5% sucrose, pH 5.7, add freshly 0.1% v/v Plant preservative mixture (APOLLO SCIENTIFIC, 26172-55-4), 25 µg/ml cefotaxime), stratified for 2 days at 4 ℃ in the dark, then transferred to growth incubator (INFORS HT Multitron) at 22 ℃ under continuous white light and 120 rpm conditions for 5 days. To induce priming, the existing medium was replaced with liquid medium containing 25 nM flg22 (primed). Control plants were handled identically but received medium lacking flg22 (naïve). The priming treatment lasted 2 h in the growth incubator, after which both primed and naïve plants were washed three times with fresh liquid medium and then grown in liquid medium for an additional 3 days. Plants were then treated with a higher dose of flg22 (100nM) as recurring stress. Both naïve and primed whole seedlings were harvested for ChIP-seq and ATAC-seq after a 3-days recovery period before the recurring stress. After recurring flg22 treatment, whole seedlings were collected for time-course RNA-seq at 5 minutes, 30 minutes, 3 hours, 24 hours and 48 hours. The transcriptional memory assay for different mutants was conducted by collecting seedlings pre-and post-30 minutes of the recurring stress.

### RNA extraction and RT-qPCR

RNA was extracted using TRIzol reagent (Invitrogen) and the Direct-Zol RNA MiniPrep (Zymo) kit, including in-column DNase I treatment following the manufacturer’s instructions. RT-qPCR analysis of *WRKY29*, *WRKY41* and *RMG1* expression in SunTag lines was performed using bulk samples of ten 7-day-old seedlings per biological replicate. Seedlings were collected immediately before treatment and 30 minutes after spraying with 100 nM flg22 (dissolved in nuclease-free H_2_O). RT-qPCRs were performed using the Luna One-Step RT-qPCR Kit (New England Biolabs, E3005E) and a CFX connect Real-time PCR detection system (Bio-Rad). IPP2 was used as the reference gene. All primers used in this study are listed in Supplementary Table 1.

### RNA-seq

For priming time-course RNA-seq, 3 biological replicates were performed for both naïve and primed Col-0 samples after recurring stress at 5 minutes (mins), 30 mins, 3 hours, 24 hours and 48 hours. Further 6 biological replicates were repeated for both naïve and primed samples immediately before (0h) recurrent stress and 30 mins after recurrent stress. Total RNA was extracted using the Direct-zol RNA miniprep kit (Zymo, R2050) including in-column DNase I treatment, with 200-500 ng of total RNA were used as input material. For the *clf-81* transcriptional memory RNA-seq experiment, two biological replicates were generated for both naïve and primed samples collected before (0h) recurrent stress in *clf-81* and Col-0. For samples collected 30 mins after recurrent stress, four biological replicates were generated for each genotype under both naïve and primed conditions. 50 ng of total RNA were used for library preparation. Libraries were prepared using QuantSeq 3’mRNA-Seq Library Prep Kit FWD (Lexogen) according to the manufacturer’s instructions and sequenced on the Illumina NovaSeq 6000 PE150 instrument. For the *jmj14* transcriptional memory RNA-seq experiment, three biological replicates were collected before (0h) recurrent stress and six biological replicates were collected 30 mins after recurrent stress for each genotype and treatment. Total RNA was submitted directly to Novogene for library preparation and sequencing.

### Machine learning

Two balanced binary classification analyses were performed using XGBoost to distinguish memory genes from either flg22-responsive non-memory genes or transcriptionally equivalent non-memory genes (10X). Features consisted of published chromatin profiles, including H3K4me1/3, H3K27me3, H2A.Z, H3K36me3, H3K56ac, H3K18ac, H3K27ac, H3K9K14ac and ATAC, which were quantified per gene. For each analysis, the respective non-memory gene set was randomly down-sampled to match the number of memory genes, and the resulting balanced dataset was split into training (80%) and test (20%) sets using stratified sampling. A fixed-parameter XGBoost model (n estimators = 100, max depth = 3, learning rate = 0.1) was trained, and five-fold stratified cross-validation was used to assess model accuracy. The final model was evaluated on the held-out test set, and SHAP values were computed on test predictions to quantify feature contributions, with summary and bar plots generated for interpretation.

### Chromatin state analysis

Chromatin states were inferred using ChromHMM (v1.25)^51,52^. Genome-wide ChIP-seq and ATAC-seq profiles were binarized into 200-bp bins using *BinarizeBam*. Models containing 2-25 states were trained using *LearnModel*, and model similarity was assessed using *CompareModels* by comparing emission profiles with the 25-state model. A 14-state model was selected as a stable and parsimonious representation of the major chromatin signatures.

Chromatin states were annotated by integrating emission probabilities, genomic-feature enrichment and associated gene-expression levels. Promoter-associated states (E1-E4) were enriched around TSSs and showed high ATAC-seq, RNA polymerase II, H3K4me3 and acetylation signals. Proximal states (E5-E7) were enriched in TSS-proximal regions and showed active chromatin features but relatively low H3K36me3. Transcription-associated states (E8-E12) were enriched across gene bodies and characterized by H3K4me1/2 and increasing H3K36me3, consistent with active transcription and elongation. E13 was dominated by H3K27me3 and annotated as Polycomb-repressed, whereas E14 showed little enrichment for the profiled chromatin features and was classified as a no-signal state. To assess chromatin-state redistribution associated with immune priming, enrichment of each state over memory-gene-associated regions was quantified separately in naïve and primed samples using *OverlapEnrichment*, enabling direct comparison of chromatin-state occupancy between the two conditions.

### Chromatin immunoprecipitation (ChIP)-seq

ChIP experiments were performed as previously described^104^. ∼1.5 g of Col-0 whole seedlings were collected from both naïve and primed samples at the end of the 3-day recovery period following the priming treatment. For the ChIPs of the SunTag:SDG2 or dSDG2:WRKY29_g1g2g3 lines, 1g 14-d-old whole plants were harvested from ½ MS plates alongside *rdr6-15* control. For all ChIPs, two or three biological replicates were performed. Tissues were ground into fine powder, and crosslinked in 25 ml Nuclear Isolation Buffer (50 mM Hepes, 1 M sucrose, 5 mM KCl, 5 mM MgCl_2_, 0.6% Triton X-100, 0.4 mM PMSF, 5 mM benzamidine, 1x Protease Inhibitor Cocktail (Roche, 11836170001)) containing 1% formaldehyde for 10 min at room temperature. Crosslinking was quenched with 0.125 M Glycine for 10 min. The homogenates were filtered through a single-layer Miracloth, and nuclei were pelleted at 2,880 × g at 4 °C for 20 min. Nuclei were resuspended with Extraction Buffer 2 (0.25 M sucrose, 10 mM Tris-HCl pH = 8.0, 10 mM MgCl_2_, 1% Triton X-100, 5 mM β-mercaptoethanol, 0.1 mM PMSF, 5 mM benzamidine, 1x Protease Inhibitor Cocktail) in 2 ml Eppendorf tube, and then centrifuged at 12,000 × g at 4 °C for 10 min. The pellets were further resuspended with Extraction Buffer 3 (1.7 M sucrose, 10 mM Tris-HCl pH = 8.0, 2 mM MgCl_2_, 0.15% Triton X-100, 5 mM β-mercaptoethanol, 0.1 mM PMSF, 5 mM benzamidine, 1x Protease Inhibitor Cocktail) and centrifuged at 12,000 × g at 4 °C for 1 h. The pellets were resuspended in 400 µl Lysis Buffer (50 mM Tris-HCl pH = 8.0, 10 mM EDTA, 1% SDS, 0.1 mM PMSF, 5 mM benzamidine, 1x Protease Inhibitor Cocktail), and diluted with 1.7 ml ChIP Dilution Buffer (1.1% Triton X-100, 1.2 mM EDTA, 16.7 mM Tris-HCl pH = 8.0, 167 mM NaCl, 0.1 mM PMSF, 5 mM benzamidine, 1x Protease Inhibitor Cocktail), and sonicated (30 s ON/30 s OFF, 21 cycles, 4°C) by Bioruptor (Diagenode). The lysate was centrifuged at maximum speed for 10 min at 4 °C, and the supernatant was transferred to fresh tubes. This clearing step was repeated twice. Chromatin was then diluted by another 2 ml ChIP Dilution Buffer. The resulting chromatin was incubated with the appropriate antibody overnight at 4 °C. The antibodies used were: anti-H3K4me1 (ab8895, Abcam), anti-H3K4me2 (ab7766, Abcam), anti-H3K4me3 (ab8580, Abcam), anti-HTA9 (AS10718, Agrisera), anti-H3K27me3 (ab6002, Abcam), anti-H3K36me3 (ab9050, Abcam). anti-H3K56ac (07-677-I, Millipore), anti-H3ac (06-599, Millipore), anti-H3K27ac (ab4729, Abcam), anti-Pol II Ser5P (ab5131, Abcam), anti-Pol II total (ab26721, Abcam), anti-HA (11867423001, Merck) and anti-H3 (ab1791, Abcam). Magnetic Protein A and Protein G Dynabeads (25 µl each, Invitrogen, 10002D and 10004D) were added and incubated for 2 h at 4 °C to capture immune complexes. Beads were washed for 5 min at 4 °C with the following buffers: Low Salt Wash Buffer (150 mM NaCl, 0.2% SDS, 0.5% Triton X-100, 2 mM EDTA, 20 mM Tris-HCl pH 8.0) twice, High Salt Wash Buffer (500 mM NaCl, 0.2% SDS, 0.5% Triton X-100, 2 mM EDTA, 20 mM Tris-HCl pH 8.0) twice, and TE buffer (10 mM Tris-HCl pH 8.0, 1 mM EDTA). Chromatin was eluted from the beads twice with 250 µl elution buffer (1% SDS, 10 mM EDTA, 0.1 M NaHCO₃) at 65 °C for 20 min each, followed by reverse crosslinking at 65 °C overnight after adding 20 µl of 5 M NaCl. Protein digestion was performed by adding 1 µl of 10 mg/ml Proteinase K, 10 µl of 0.5 M EDTA (pH 8.0), and 20 µl of 1 M Tris-HCl (pH 6.5), and incubating the mixture at 45 °C for 4 h. DNA was extracted with 550 µl phenol-chloroform-isoamyl alcohol (25:24:1, Sigma, P3803) followed by 550 µl chloroform. The aqueous phase was precipitated with 50 µl 3 M sodium acetate (Thermo, 10190890), 2 µl GlycoBlue (Invitrogen, AM9516), and 1 ml 100% ethanol at -20 °C overnight. DNA was pelleted by centrifugation at maximum speed for 1 h at 4 °C, washed with 1 ml 70% ethanol, air-dried, and resuspended in nuclease-free water. The ChIP-purified DNA was directly used for ChIP-qPCR and ChIP-seq library preparation. ChIP libraries were generated using NuGen Ovation Ultra Low System V2 kits according to the manufacturer’s instructions. The libraries were sequenced for PE150 reads on an Illumina NovaSeq X instrument.

### Sequential ChIP-seq

Tissue fixation, chromatin sonication, and immunoprecipitation with the first antibody followed the procedures described above, except for specific modifications noted below. For each replicate, 4 g of 14-day-old Col-0 seedlings grown on ½ MS plates were used. After quenching, the chromatin lysate was filtered first through a single layer of Miracloth and subsequently through a double-layer filter. Before adding the antibody into ChIP Dilution Buffer diluted chromatin, the chromatin was pre-cleared by magnetic Protein A and Protein G Dynabeads (15 µl each) at 4 °C for 2 h. After first antibody (H3K27me3) immunoprecipitation, the beads were washed by Low salt buffer (2x), High salt buffer (2x), LiCl (250 mM LiCl, 1% Igepal, 1% sodium deoxycholate, 1 mM EDTA, 10 mM Tris-HCl pH = 8.0, 2x) and TE buffer (2x). The protein-DNA complexes were eluted from beads using reChIP elution buffer (1× TE, 2% SDS, 15 mM DTT, 1x Protease Inhibitor Cocktail) for 15 min at 37 °C at 400rpm. The supernatant was collected and diluted to a final volume of 400 µl with ChIP Dilution Buffer supplemented with 50 µg BSA and protease inhibitors. The chromatin was then divided into four 100 µl aliquots: one reserved as the first-IP input, one used as a second-IP positive control with H3K27me3 antibody, one for H3K4me3 immunoprecipitation, and one processed as a no-antibody control. Each 100 µl aliquot was diluted to 1 ml before the second immunoprecipitation. The second IP was then performed by incubating the samples overnight at 4 °C. After binding to Dynabeads, the beads were washed sequentially with low-salt buffer (3×), high-salt buffer (3x), LiCl buffer (2x), and TE buffer (2x), followed by elution and DNA purification as described in the standard ChIP protocol. The libraries were generated using NuGen Ovation Ultra Low System V2 kits according to the manufacturer’s instructions. The libraries were sequenced for PE150 reads on an Illumina NovaSeq X instrument.

### Assay for Transposase-Accessible Chromatin (ATAC)-seq

ATAC experiments were performed based on the protocol previously described^105^ with minor modifications. Briefly, ∼80 Col-0 whole seedlings were collected from both naïve and primed samples at the end of the 3-day recovery period following the priming treatment. Two biological replicates were performed. Tissues were fixed in Fixative Buffer (1× PBS containing 1% formaldehyde) and quenched in Glycine Buffer (0.125 M Glycine in PBS) using rapid vacuum infiltration with a 10 ml syringe. Nuclei were released by gently chopping the tissue for 5 min in a Petri dish using a sterile razor blade in 1,000 µl of General purpose buffer (0.5 mM spermine·4HCl, 30 mM sodium citrate, 20 mM MOPS, 80 mM KCl, 20 mM NaCl, 0.5% Triton X-100, pH 7.0). The resulting slurry was passed sequentially through cell strainers with decreasing pore sizes (70 µm, 40 µm, and 35 µm) and collected into a Corning Falcon tube. Nuclei were sorted on a SAFB FACS Aria III high-speed cell sorter. Prior to sorting, 1 µl of Hoechst stain was added to each sample. Nuclei were then sorted using a 70 µm cytonozzle with PBS (pH 7.0) as the sheath fluid at 30.5/30.0 psi (sample/sheath), using a 1-drop purity mask and triggering on Hoechst emission (450/50 nm). All Hoechst-positive nuclei populations-including 2C, 4C, 8C, and higher-ploidy fractions-were collected to ensure comprehensive chromatin representation across endoreduplicated states. The 2C population was defined as the first distinct Hoechst-emitting peak clearly separated from background scatter. Approximately 50,000 nuclei were collected per sample per replicate. Prior to tagmentation, sorted nuclei were incubated at 60 °C for 5 min. Nuclei pellets were then treated with 1 µl Tn5 transposase (Illumina, 20034198) at 37 °C for 40 min. The reaction volume was brought to 200 µl with SDS buffer (50 mM Tris-HCl pH 8.0, 1% SDS, 10 mM EDTA pH 8.0), followed by the addition of 8 µl 5 M NaCl for de-crosslinking overnight at 65 °C. DNA was purified using a DNA Clean & Concentrator kit (Zymo, D4013). 3 ul of purified DNA was used for library preparation using Q5 polymerase and indexed i5 and i7 primers (Nextera, 53155). Libraries were purified with AMPure beads (Beckman, A63881) and sequenced for PE150 reads on an Illumina NovaSeq X instrument.

### snRNA-seq

Nuclei were isolated from the shoots of Arabidopsis seedlings grown vertically on MS plates for 11 days, 100 seedlings per treatment. Seedlings were spray-treated with either mock (Nuclease free H_2_O (nfH_2_O)) or 100 nM flg22 (CAT: crb1000331j dissolved in nfH_2_O) treatment for 30 minutes prior to separation of the roots and the shoots using a razor blade. Nuclei were then extracted from shoot tissue following a previously published protocol with slight adaptation^68^. Briefly, shoots were chopped for 2 minutes using a razor blade in a 5-cm Petri dish containing 2ml of freshly prepared nuclei isolation buffer (NIB: 10mM Tris-HCL pH7.4, 10mM NaCl, 3mM MgCl_2_, 0.5mM spermidine, 0.2mM spermine, 1x Roche complete protease inhibitors (EDTA free), 0.01% Triton-X, 1% BSA and 1 U μl^−1^ Protector RNase Inhibitor). Solution was filtered through a 70-μm and then through a 30-μm cell strainer. Nuclei were presorted using an Aria™ III Cell Sorter with a 70-μm nozzle, and due to being too concentrated, diluted with extra 2 ml of NIB. Nuclei were then stained with 2 μl Hoechst solution (Hoechst 33342, trihydrochloride, trihydrate; 10 mg ml^−1^ solution in water, Invitrogen) and sorted. Sorted nuclei were collected in a 1.5-ml tube containing 100 μl of catch buffer (10mM Tris-HCL pH7.4, 10mM NaCl, 3mM MgCl_2_, 0.5% BSA and 1 U μl^−1^ Protector RNase Inhibitor). Final volume after sort was ∼200 μl. 10 μl were moved to a hemocytometer for quantification (Nano Entek C-CHIP DHC – N01) under a light microscope. Concentrations were between 400 and 500 nuclei/μl per sample. 2 biological replicates were used per treatment with each replicate consisting of 10000 nuclei. We then proceeded to generate cDNA, barcode and amplify and construct (dual index) the library using a Chromium Next GEM Single Cell 3′ Reagent Kits v.3.1 following the manufacturer’s protocol. Libraries were sequenced using Illumina NovaSeq.

### *Pseudomonas syringae* bioluminescence infection assay (*Pst::LUX*)

The *Pst::LUX* colonisation assay was performed as previously described^53^. In brief, seven-day-old Arabidopsis seedlings grown in 96-well plates on 1/2× MS medium supplemented with 200 μg/ml Timentin were spray-inoculated with bioluminescent *Pseudomonas syringae pv.* tomato DC3000::LUX (OD_600_ = 0.2 in 10 mM MgSO₄ containing 0.015% Silwet L-77). Plates were sealed with parafilm to maintain high humidity and imaged for 4 subsequent days. Prior to imaging, plates were dark-adapted for 2 min and imaged using an ImageQuant 800 system with a 4-minute exposure time. Corresponding brightfield images were acquired for well localization. Bioluminescence was quantified in Fiji by applying well outlines from the brightfield images to the luminescence images and extracting the mean signal intensity per well.

## Bioinformatic analysis

### Sources of published data

Previously published RNA-seq data (E-MTAB-9694)^33^ were incorporated to enable direct comparison with the datasets generated in this study. ChIP-seq data of H3 (SRX5534451), H2A.Z (SRX5534457), H3K4me1 (SRX5534456), H3K4me3 (SRX5534455), H3K27me3 (SRX5534452), H3K36me3 (SRX5534453), H3K56ac (SRX5534454) and ATAC-seq (SRR8742423, SRR8742424, SRR8742425)^106^; H3K27ac, H3K9K14ac (E-MTAB-7611 and E-MTAB-8265)^107^; H3K18ac (SRX2533656)^108^ were used as the features for Machine learning and conventional ChIP-seq overlapping peaks in Fig. 3. ATAC-seq datasets (GSE128434)^106^ and (GSM2897831)^109^ were used for direct comparison with the datasets generated in this study. Published H3K4me3 ChIP-seq data from the *jmj14* mutant and JMJ14-FLAG ChIP-seq data (GSE204681)^47^, used to assess JMJ14 occupancy, were re-analyzed in this study.

### RNA-seq

In accordance with the manufacturer’s (Lexogen) instructions, only the read (*1.fq.gz) from PE150 bp data was used for downstream analysis. Briefly, reads were trimmed using cutadapt (version 1.18) and were mapped to the genome using STAR (version 2.7.10b) to the TAIR10 genome (--quantMode GeneCounts --alignIntronMax 10000 -- outSAMmultNmax 20). Gene counts were used as input for DESeq2 analysis in R. Differentially expressed genes were defined using an adjusted p-value cutoff of <0.01. Motifs identification was performed using HOMER^110^ (findMotifsGenome.pl). For the unbiased clustering the time-course data, kallisto^111^ extracted TPMs were used as input for analyses using the Clust^112^ package in R, with default parameters used to obtain the 15 clusters analysed. Gene ontology (GO) analyses were performed using the topGO package in R.

### ChIP-seq, Sequential ChIP-seq and ATAC-seq

Both the previously published datasets and those generated in this study (ChIP-seq, Sequential ChIP-seq and ATAC-seq) were analysed as described below. The reads in fastq format were aligned to the TAIR10 reference genome (--no-unal) using Bowtie2 (version 2.5.4) and was converted to bam format using Samtools (version 1.22.1). Reads were de-duplicated using the Samtools fixmate and markdup commands. Deeptools (version 3.5.5) was used to generate the tracks using bamCoverage (-- normalizeUsing RPGC, --effectiveGenomeSize 135000000 --binSize 10) with multicopy regions blacklisted (--blackListFileName) using the regions identified in ref^113^. Peaks were called using MACS2 (version 2.2.9.1). For published data, parameters (-q 0.1) and (--broad --broad-cutoff 0.05) were used for narrow peaks (H3K27ac, H3K9K14ac, H3K18ac, H3K4me3, H2A.Z, H3K56ac) and broad peaks (H3K4me1, H3K27me3, H3K36me3), respectively. Parameters (-q 0.01) and (-p 0.01) were used for narrow peaks in ChIP-seq and Sequential ChIP-seq, respectively. For ATAC-seq, peaks called with default parameters. Peaks were first called for each biological replicate and subsequently merged using bedtools intersect (version 2.31.1) to identify reproducible peaks shared across replicates. Peaks were annotated by ChIPseeker in R (version 4.4.1). Correlation plots were generated in DeepTools using MultiBamSummary or MultiBigwigSummary (--binSize 100) and PlotCorrelation (-c pearson/spearman --skipZeros --removeOutliers --plotNumbers -p heatmap). The IGV genome browser was used to visualize the data. Deeptools was used to generate the metaplots and heatmaps.

### snRNA-seq

#### Data processing and quality control

Sequencing reads were processed with CellRanger V.7.0.0 (10X genomics) using default parameters to generate count matrices aligned to the Arabidopsis thaliana reference genome (TAIR10). Downstream analysis was performed in R using Seurat (v.5.3.0). For each sample, a Seurat object was created from the filtered feature-barcode matrix and standard quality control metrics were computed per nucleus, including the number of detected genes, total unique molecular identifier (UMI) counts and the fraction of reads mapping to mitochondrial genes. Low-quality nuclei were removed by retaining only nuclei with more than 200 detected genes and < 5% mitochondrial genes. Each dataset was normalized independently using Seurat’s NormalizeData function (log-normalization with default parameters).

### Data integration, dimensionality reduction and clustering

Batch correction across biological replicates and treatments was performed using Seurat’s anchor-based integration workflow. Integration anchors were identified with FindIntegrationAnchors(), and the data were integrated with IntegrateData(). A shared nearest-neighbor graph was constructed using FindNeighbors(), and unsupervised clustering was carried out with FindClusters(). Dimensionality reduction for visualization was performed using RunUMAP(). Cell type annotation was carried out after clustering by comparing cluster-enriched marker genes with established *Arabidopsis thaliana* shoot and leaf marker genes obtained from a curated reference dataset^36^. Cluster markers identified by FindAllMarkers() were overlapped with this marker list, and annotation was supported by UCell signature scores calculated for each curated marker set. Expression of canonical markers representing major leaf cell types (including epidermis, guard cells, trichomes, mesophyll, phloem companion cells and bundle sheath cells) to confirm and refine cluster identities. Gene Ontology enrichment analyses were performed using clusterProfiler (4.14.6) with org.At.tair.db as the annotation database and TAIR identifiers as the key type. Analyses were restricted to Biological Process (BP) terms. *P*-values were adjusted using the Benjamini–Hochberg method, with significance thresholds of p < 0.05 and q < 0.2.

For the Wang *et al.,* (zenodo.16533111)^38^ and Nobori *et al.,* (GSE226826)^37^ datasets, both objects were downloaded as processed SeuratObject (v5.4.0) using author specified annotations. Cell-type restriction was quantified for each gene as the squared coefficient of variation (CV^2^) of pseudobulk CPM across the 23 RNA clusters from Nobori et al and the 30 clusters from Wang et al. To control for expression levels between the memory gene set and non-memory responsive genes, the 1,414 flg22-responsive non-memory genes were divided into 20 pseudobulk CPM bins of equal size, and each of the 2,000 permutations (tidyverse in R (v4.5.3)) drew a control set of the same size as the memory gene set with the same number of genes per bin.

## Data availability

All sequencing data have been submitted to the NCBI Gene Expression Omnibus (GEO) under accession numbers GSE346174 and GSE346373.

## Code availability

Custom scripts used are available on GitHub at https://github.com/C-Jake-Harris/

## ACKNOWLEDGEMENTS

We thank Dr. Matthew Naish, Dr. Régis Lopes Corrêa, and Prof. Israel Ausin for kindly providing seeds. We would like to thank Leonie Luginbühl and Tianshu Sun for their invaluable assistance with the single nuclei RNA-seq transcriptomics. We would also like to thank Jurriaan Ton and Adam Parker for their help in setting up the *Pst::LUX* experimental pipeline in our group. C.J.H. is supported by a Royal Society University Research Fellowship (URF\R1\201016) and an ERC Starting Grant (TransPlantMemory), funded by UKRI EPSRC (EP/X025306/1). J.B. is supported by a Marie Curie postdoctoral fellowship, funded by UKRI (EP/Z001749/1). H.T. is supported by a 4-year BBSRC-funded doctoral studentship. V.A. is supported by a 4-year BBSRC-funded doctoral studentship.

## AUTHOR CONTRIBUTIONS

C.J.H. conceived, designed and supervised the research. C.J.H. performed time course RNA-seq, NGS bioinformatic analysis. L.X. performed sequential ChIP-seq, ChIP-seq, NGS bioinformatic analysis and mutants screening. J.B. performed snRNA-seq, NGS bioinformatic analysis. L.X. and J.B. performed the experiments relating to SunTag lines. B.W. performed the experiments relating to *clf* mutant. L.X. and H.T. performed ATAC-seq. V.A. performed experiments validating *bona fide* priming activity. C.J.H., L.X. and J.B. wrote the manuscript with the participation of all the other authors. All authors read and approved the final manuscript.

---

**Supplementary Table 1.**
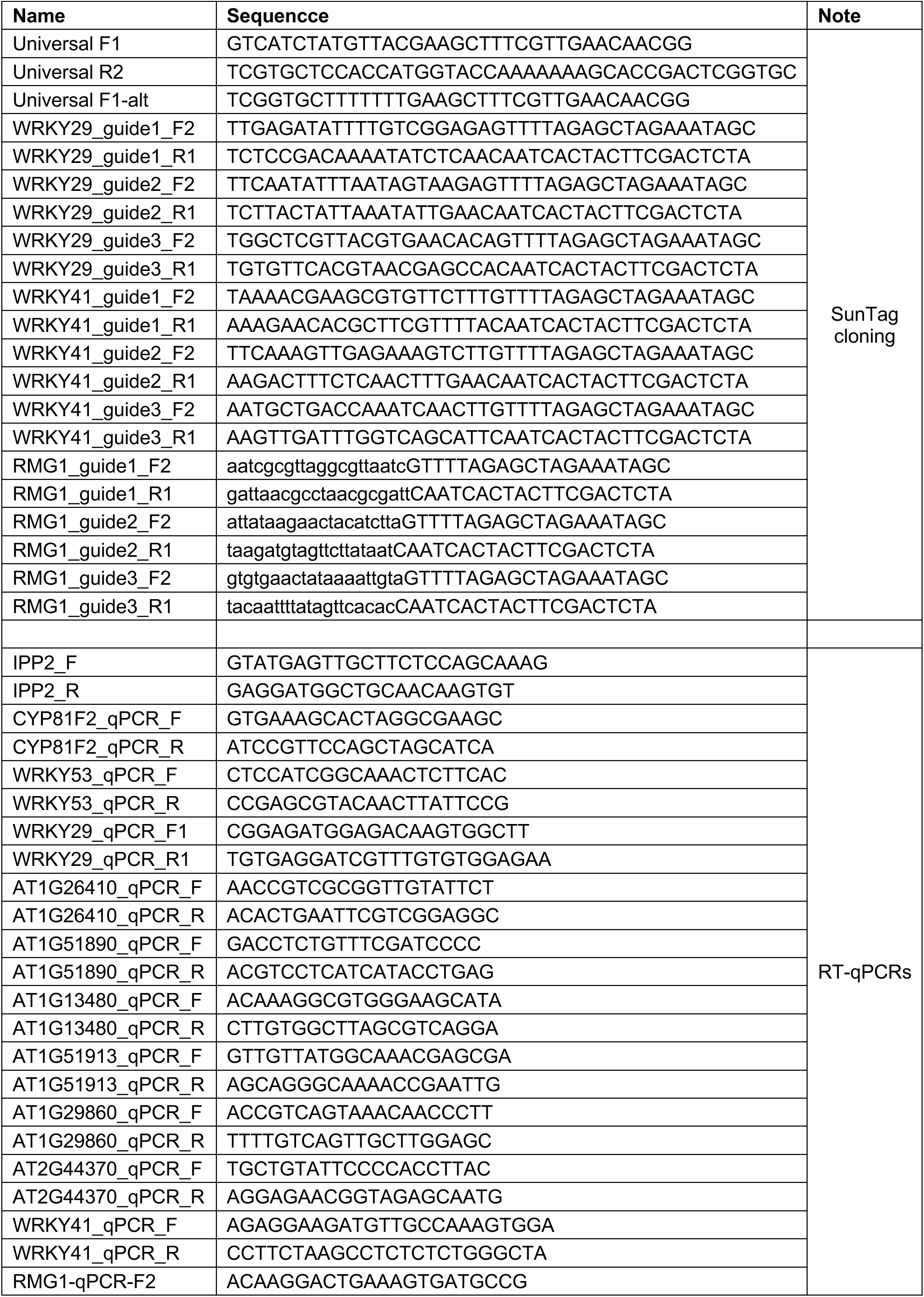
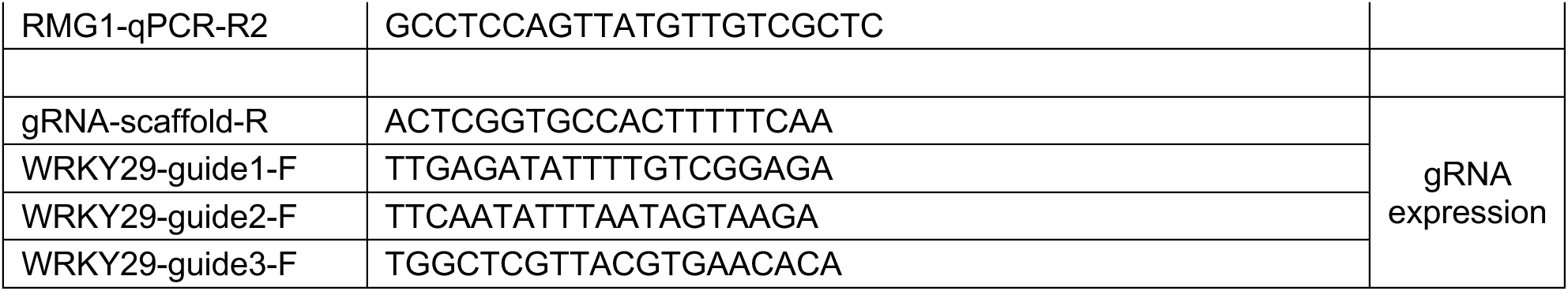
: Oligo list.

**S Fig. 1.**
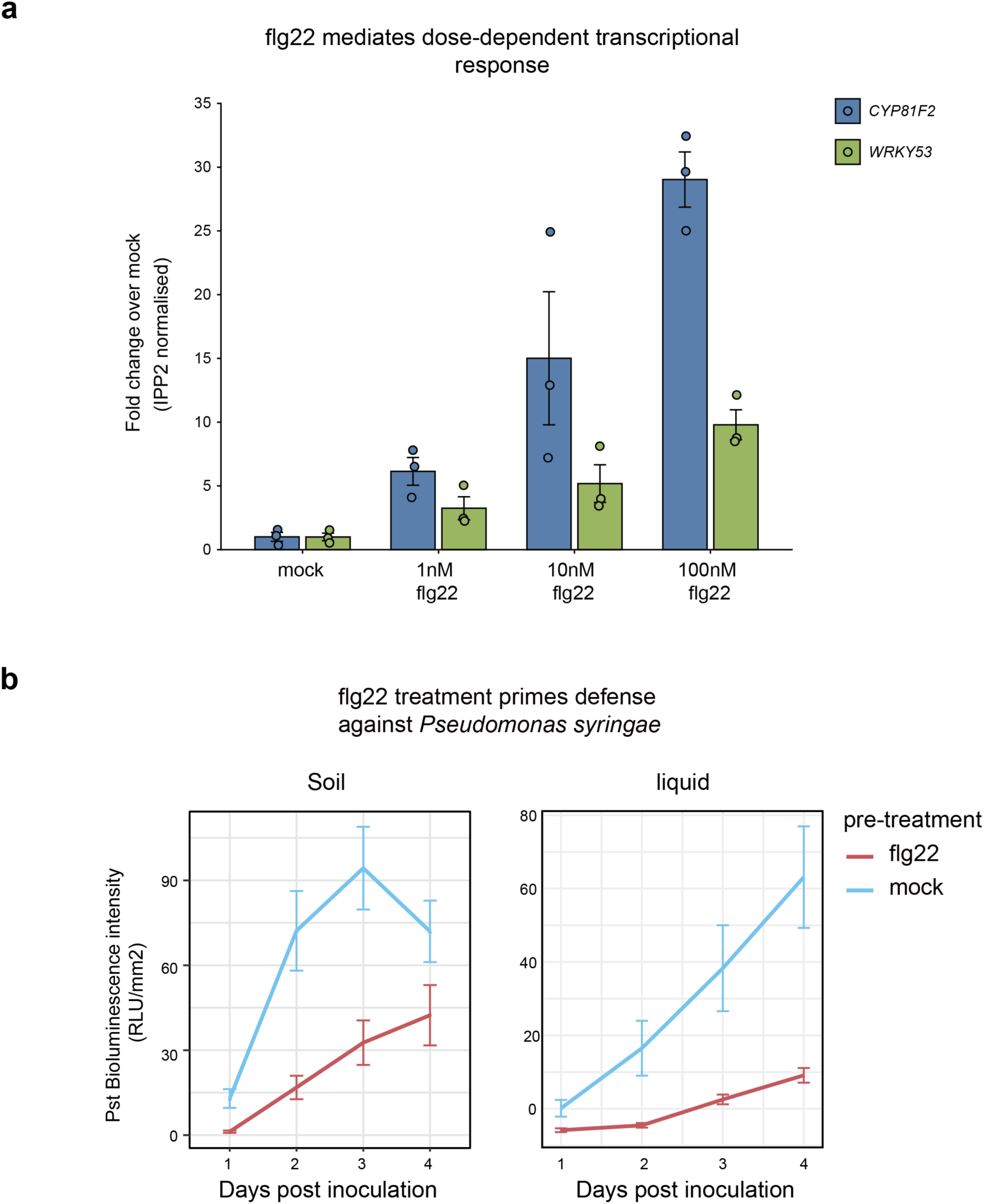
Flg22-mediated transcriptional responses and defense priming. a. Bar plot showing the expression of the flg22-responsive genes *CYP81F2* and *WRKY53* following treatment with mock and 1 nM, 10 nM, or 100 nM flg22. Transcript levels were normalized to IPP2 and are shown as fold change relative to mock-treated samples. Bars represent mean ± SEM, with individual biological replicates shown as points (n = 3 biological replicates per treatment). **b.** *Pst*::LUX bioluminescence assay quantifying bacterial colonization at 1, 2, 3, and 4 days post-inoculation in plants pretreated with flg22 or mock under soil-and liquid-grown conditions. Bioluminescence intensity is shown as relative light units (RLU/mm²). Data are presented as mean ± SEM.

**S Fig. 2.**
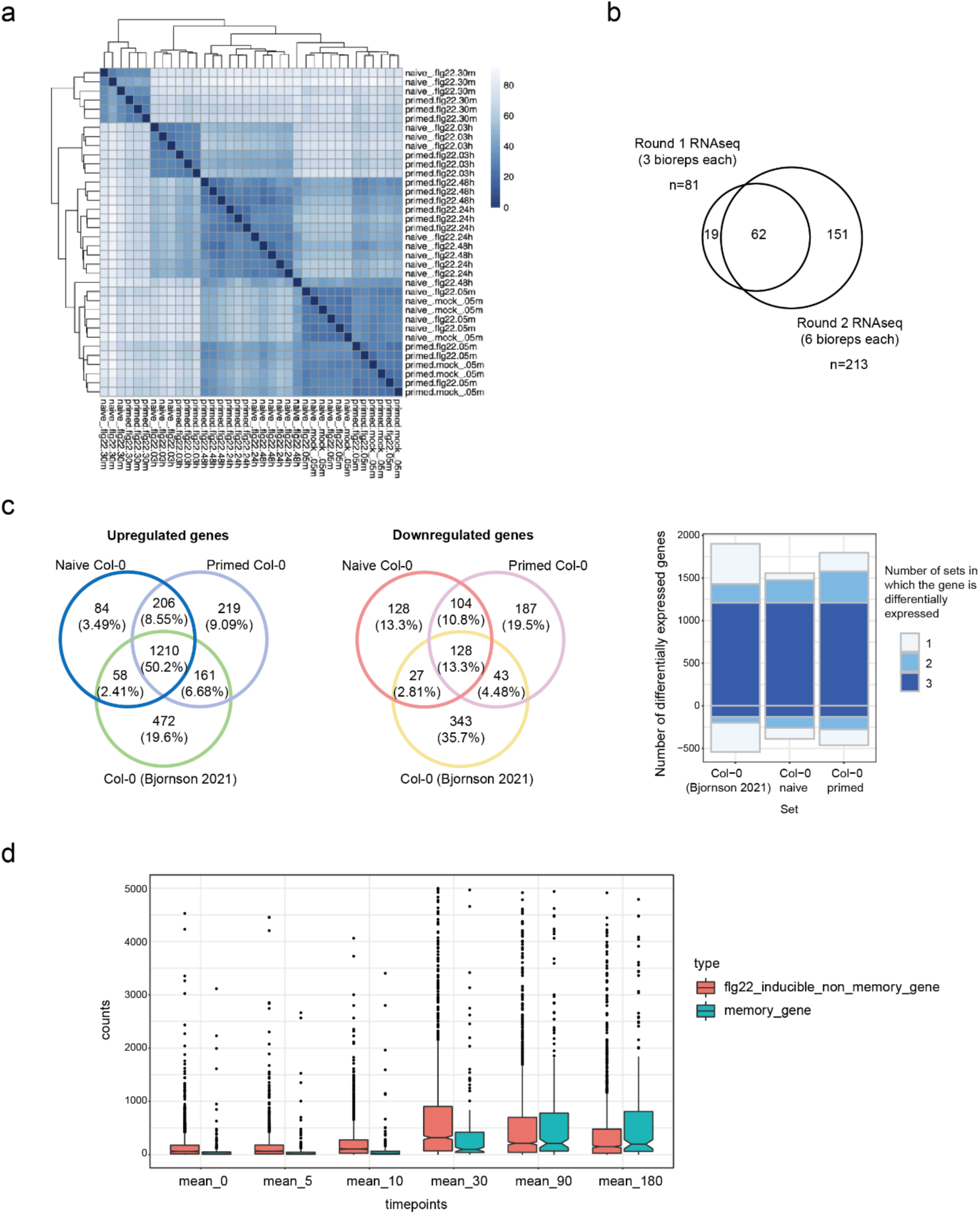
Quality control of the time-course RNA-seq.a. Pairwise correlation analysis of the time-course RNA-seq expression profiles from naïve and primed seedlings treated with flg22 or mock at 5 minutes (mins), 30 mins, 3 hours, 24 hours, and 48 hours after the second treatment (n = 3 biological replicates per condition and time point). Hierarchical clustering reflects the similarity between samples across conditions and time points **b.** Venn diagram showing the overlap of memory genes identified at 30 mins after recurrent flg22 treatment in round 1 RNA-seq (3 biological replicates) and round 2 RNA-seq (6 biological replicates). **c.** Venn diagrams showing the overlap of upregulated genes (left) and downregulated genes (middle) among naïve Col-0, primed Col-0, and Col-0 from Bjornson *et al*. (2021). Right, stacked bar plot showing the number of differentially expressed genes grouped by the number of datasets in which each gene was identified. **d.** Boxplots showing the distribution of RNA-seq counts from Bjornson *et al*. (2021) for memory genes and flg22-inducible non-memory genes across the indicated time points. Boxes indicate the interquartile range, with the median shown as a horizontal line; whiskers denote 1.5x the interquartile range (IQR), and dots represent individual genes.

**S Fig. 3.**
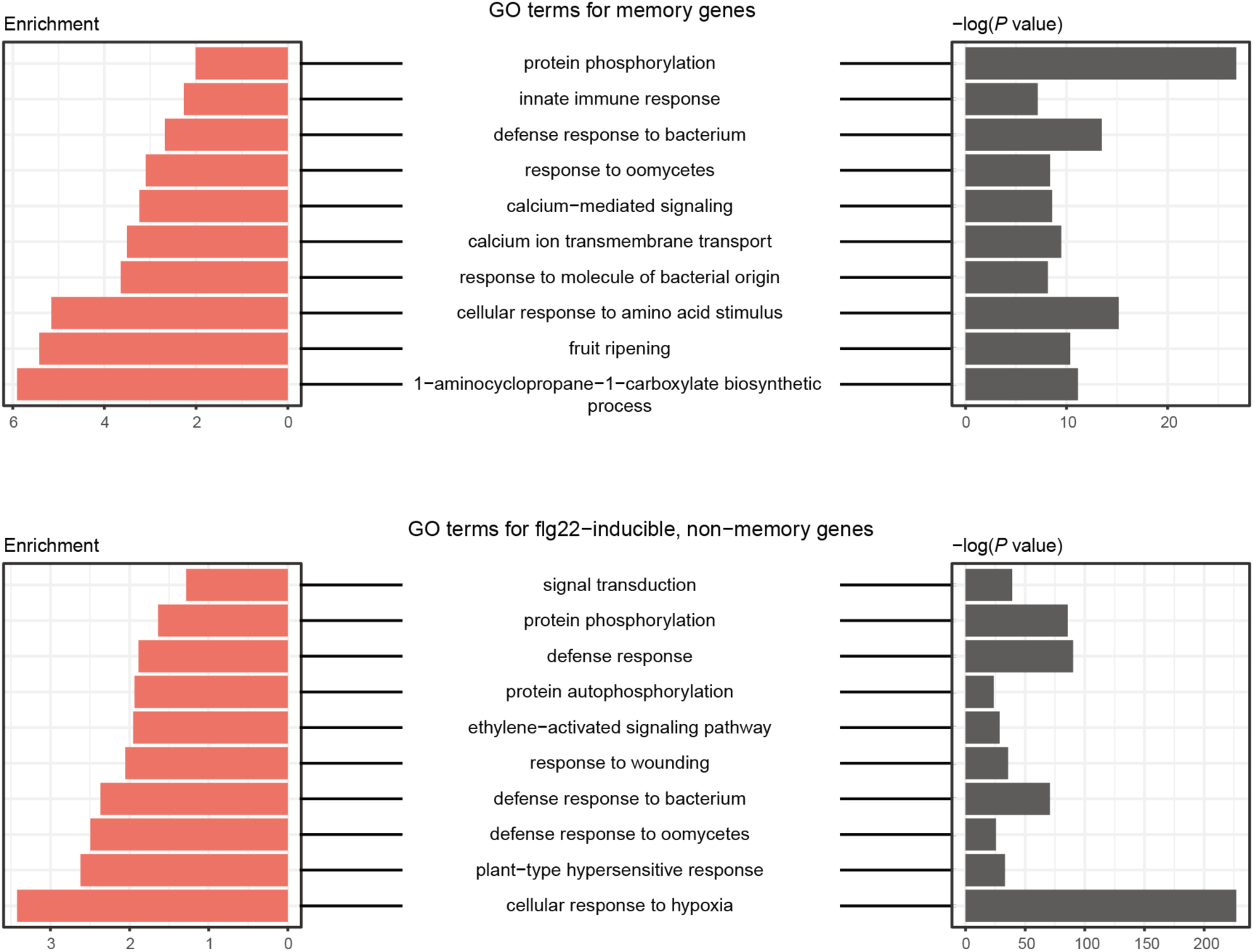
Gene Ontology (GO) term analysis of memory genes (upper panel) and flg22-inducible non-memory genes (lower panel).

**S Fig. 4.**
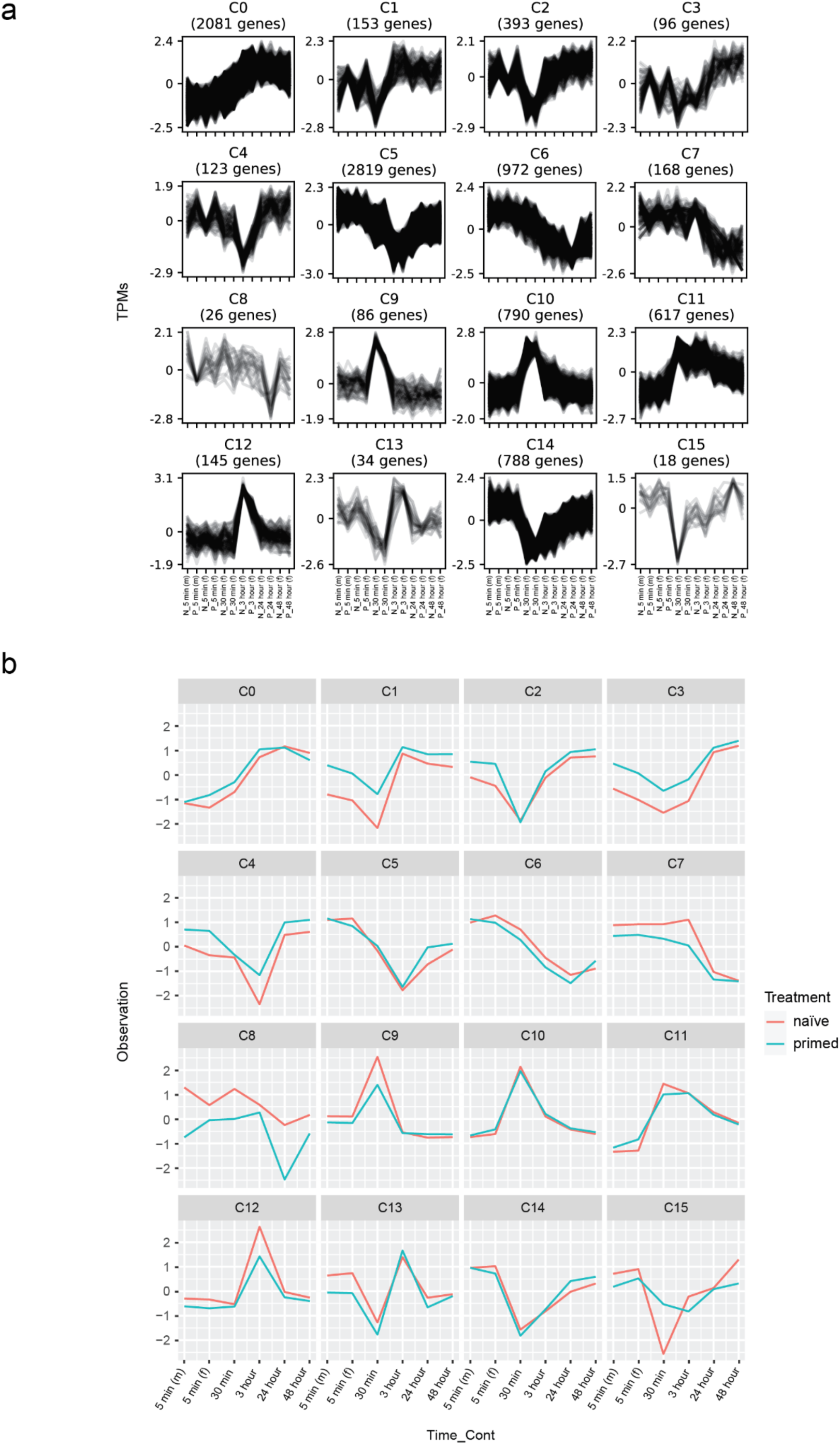
Time-course clustering of gene expression during naïve and primed flg22 responses. a. Genes were grouped into 16 clusters (C0-C15) based on their expression dynamics. Each panel shows normalized transcript abundance (TPM) over the time course for both naïve (N) and primed (P) samples. The number of genes in each cluster is indicated. **b.** Average expression profiles of genes within each cluster (C0-C15), comparing naïve and primed samples across the time course. Lines represent the mean normalized expression for each cluster under the indicated treatment conditions.

**S Fig. 5.**
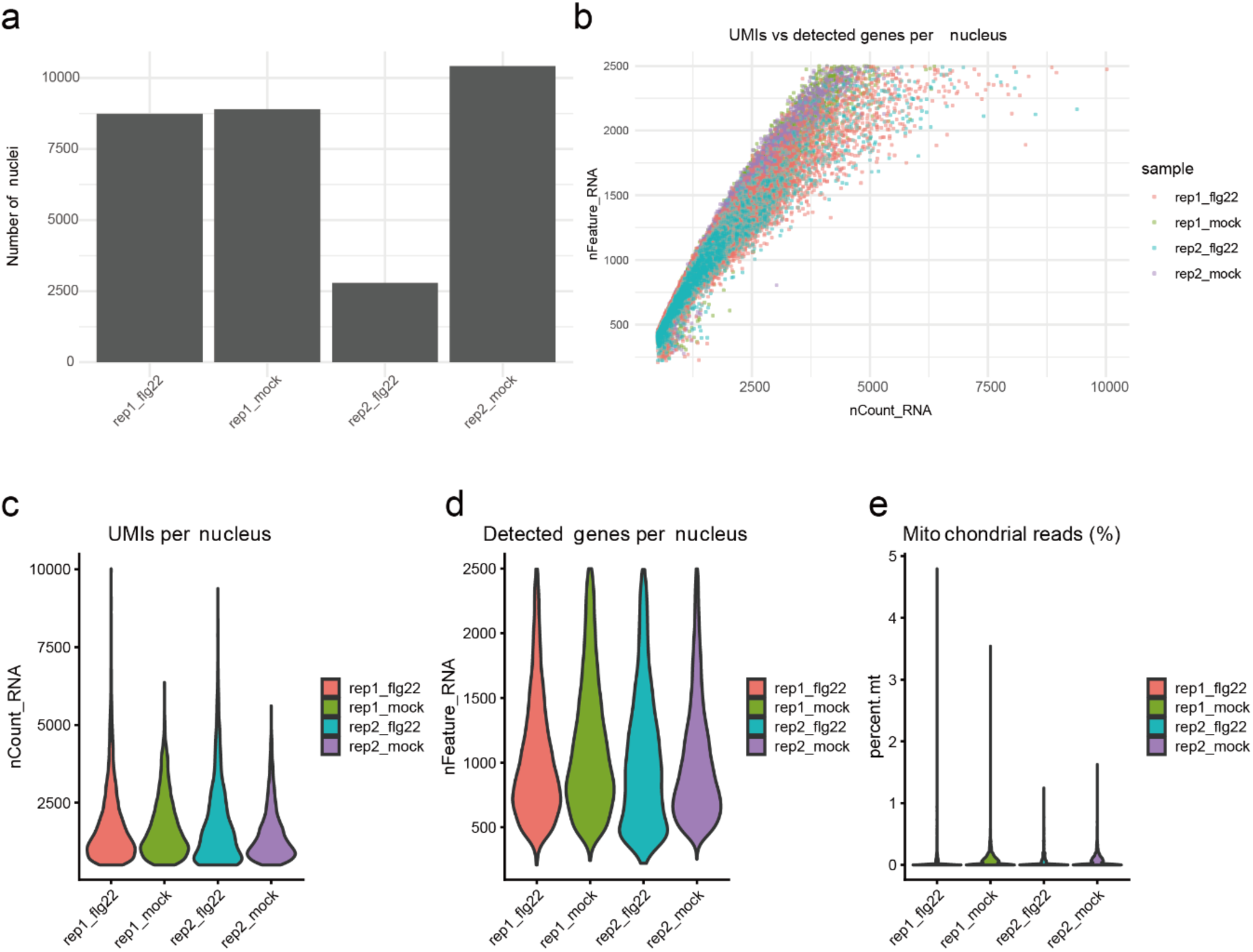
snRNA-seq quality control. a. Bar plot showing the number of nuclei retained in each snRNA-seq sample after quality control filtering. Nuclei were isolated from shoot tissue collected 30 mins after flg22 or mock treatment, with two biological replicates per treatment. **b.** Scatter plot showing the relationship between the number of detected genes per nucleus (nFeature_RNA) and total UMI counts (nCount_RNA) across all nuclei. Each point represents a single nucleus, colored by sample. **c.** Violin plots showing the distribution of total UMI counts per nucleus (nCount_RNA) across the four snRNA-seq samples. **d.** Violin plots showing the distribution of the number of detected genes per nucleus (nFeature_RNA) across the four snRNA-seq samples. **e.** Violin plots showing the distribution of mitochondrial read percentages per nucleus across the four snRNA-seq samples.

**S Fig. 6.**
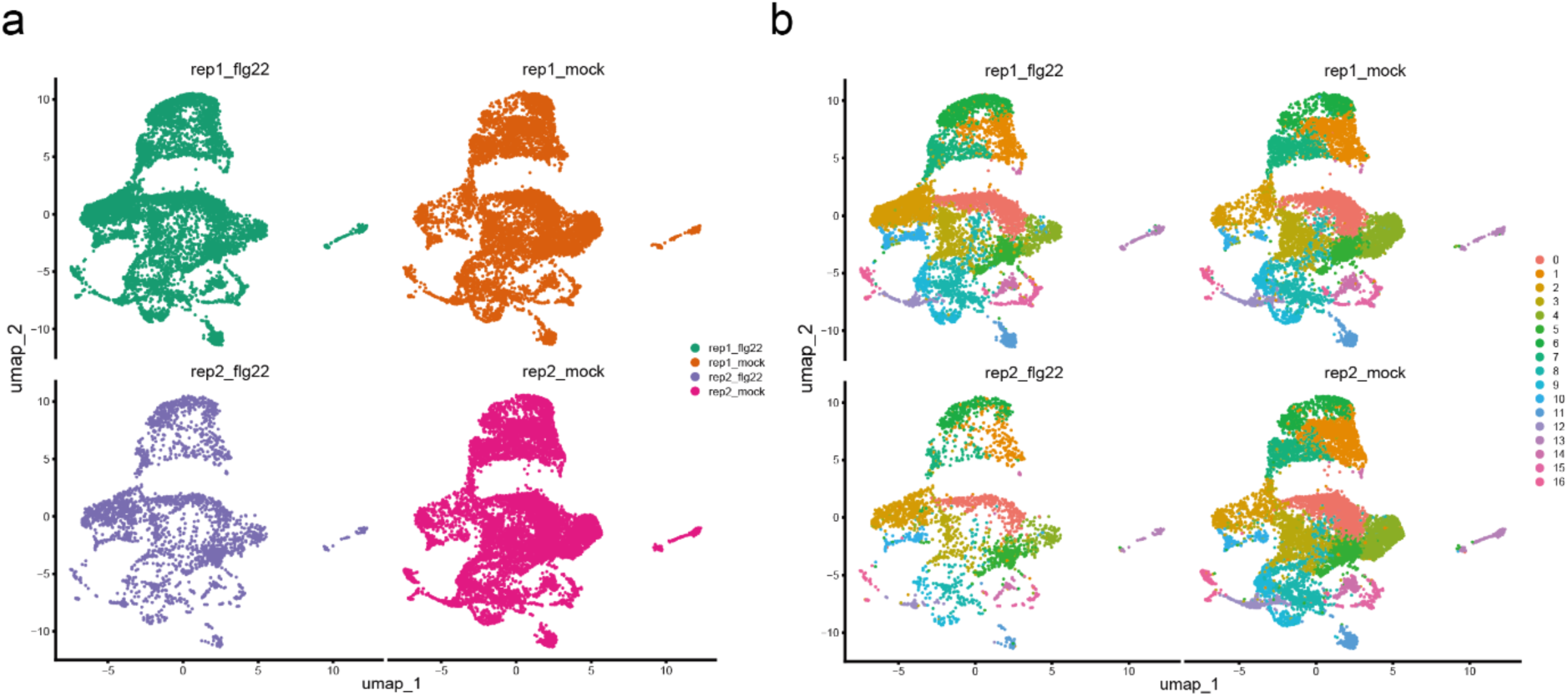
Replicate integration and cluster consistency of snRNA-seq. a. UMAP embeddings of nuclei from shoot tissue collected 30 mins after flg22 or mock treatment, shown separately for each snRNA-seq library. Two biological replicates were included for each treatment. **b.** UMAP visualization of the integrated snRNA-seq dataset coloured by cluster identity and shown separately for each sample. Nuclei were assigned to 17 transcriptional clusters (0-16). The comparable distribution of clusters across biological replicates and treatments indicates that the major cell populations were consistently represented across samples.

**S Fig. 7.**
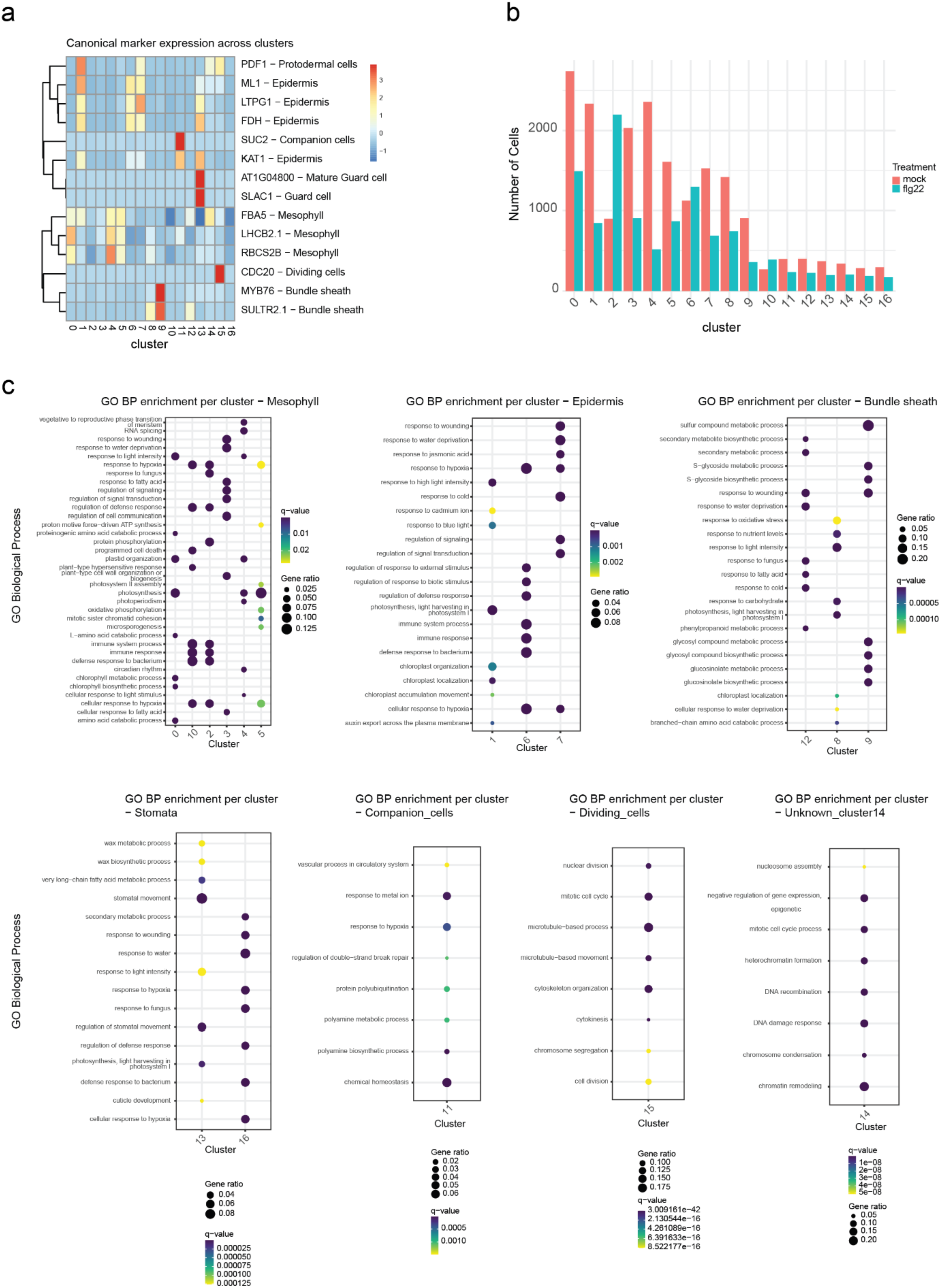
Cell-type annotation and functional characterization of snRNA-seq clusters. a. Heatmap showing the expression of canonical cell-type marker genes across snRNA-seq clusters. Marker genes used for annotation are indicated together with their corresponding cell types, including protodermal, epidermal, companion, guard, mesophyll, dividing, and bundle sheath cells. Expression values are scaled across clusters. **b.** Bar plot showing the number of nuclei assigned to each cluster in mock-and flg22-treated samples. Nuclei were isolated from shoot tissue collected 30 mins after treatment, with two biological replicates per treatment combined for visualization. **c.** Gene Ontology (GO) Biological Process enrichment analysis for genes associated with the indicated cell-type clusters. Dot size represents the gene ratio, and colour represents the enrichment significance (*q*-value). Clusters are grouped according to their annotated cell types, including mesophyll, epidermis, bundle sheath, stomatal, companion, dividing, and unassigned clusters.

**S Fig. 8.**
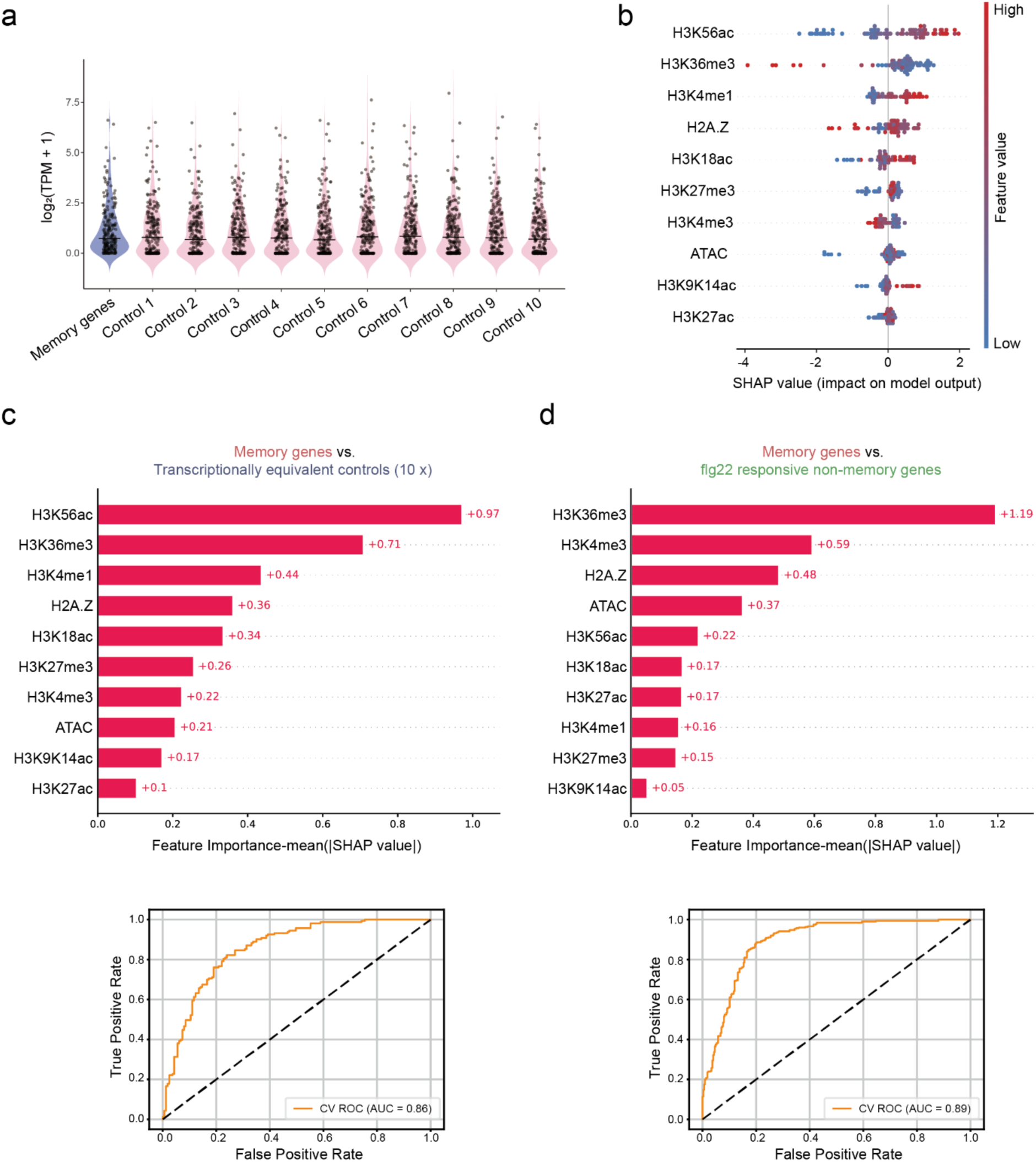
Chromatin features associated with memory genes. a. Violin plots showing the distribution of gene expression levels (log_2_(TPM + 1)) for memory genes and 10 transcriptionally equivalent control gene sets (Control 1 - Control 10). This analysis serves as a sanity check to confirm comparable expression distributions between memory genes and the control gene sets. Each dot represents an individual gene. **b.** SHAP summary plot showing the contribution of chromatin features to XGBoost models trained to distinguish memory genes from 10 groups of transcriptionally equivalent control genes. Features are ranked by importance, with SHAP values indicating their impact on model output and point colour representing the relative feature value. **c.** Bar plot showing the mean feature importance, measured by absolute SHAP values, for chromatin marks and accessibility features in XGBoost models distinguishing memory genes from 10 transcriptionally equivalent control gene sets. Bottom, receiver operating characteristic (ROC) curve showing model performance (AUC = 0.86). **d.** Bar plot showing the mean feature importance, measured by absolute SHAP values, for chromatin marks and accessibility features in an XGBoost model distinguishing memory gene from flg22-responsive non-memory genes. Bottom, receiver operating characteristic (ROC) curve showing model performance (AUC = 0.89).

**S Fig. 9.**
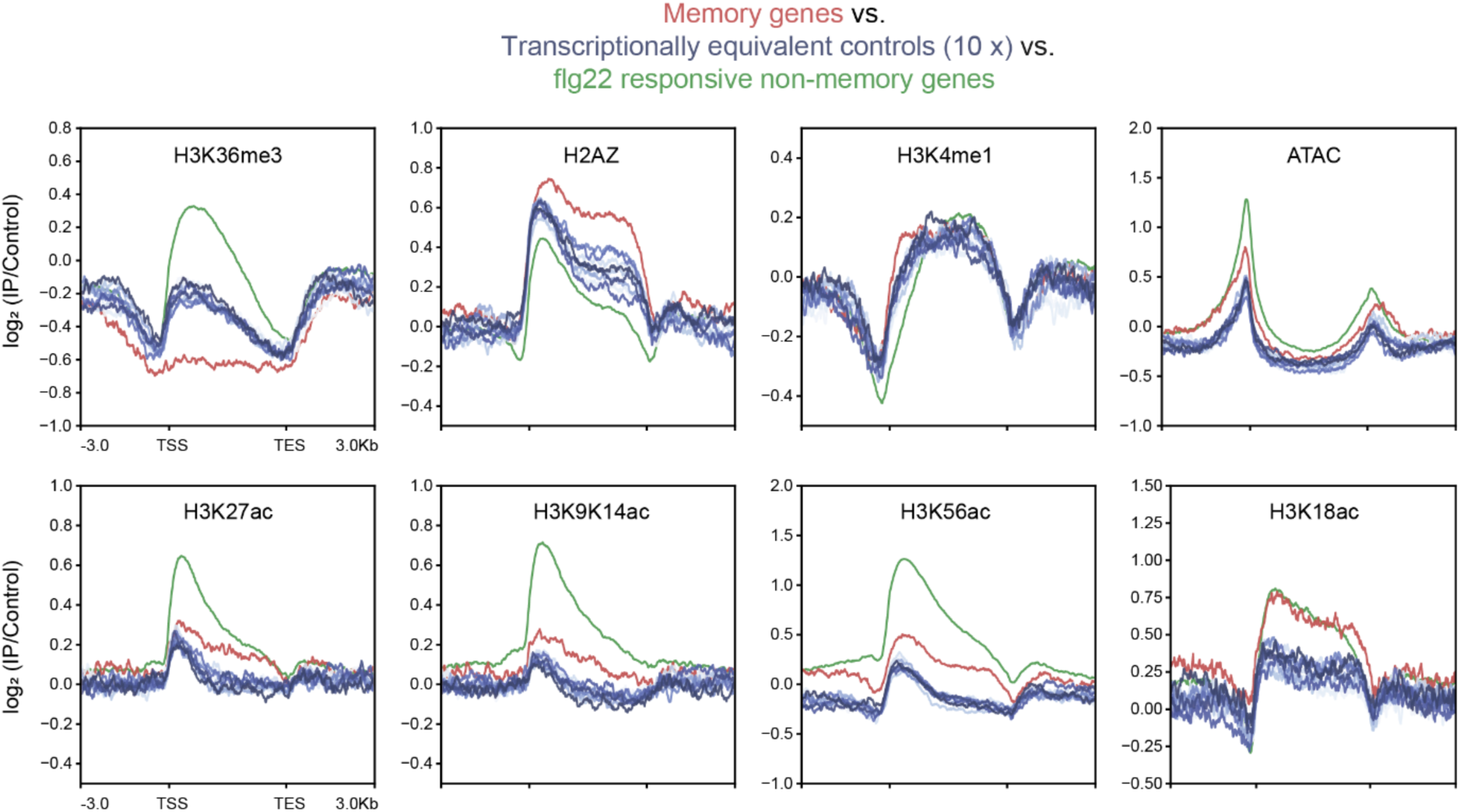
Chromatin profiles of memory genes compared with transcriptionally equivalent controls and flg22-responsive non-memory genes. Metaplots showing H3-or input-normalized chromatin profiles across memory genes (red), ten sets of transcriptionally equivalent control genes (blue), and flg22-responsive non-memory genes (green). Profiles are shown for H3K36me3, H2A.Z, H3K4me1, H3K27ac, H3K9K14ac, H3K56ac, and H3K18ac, together with ATAC-seq accessibility.

**S Fig. 10.**
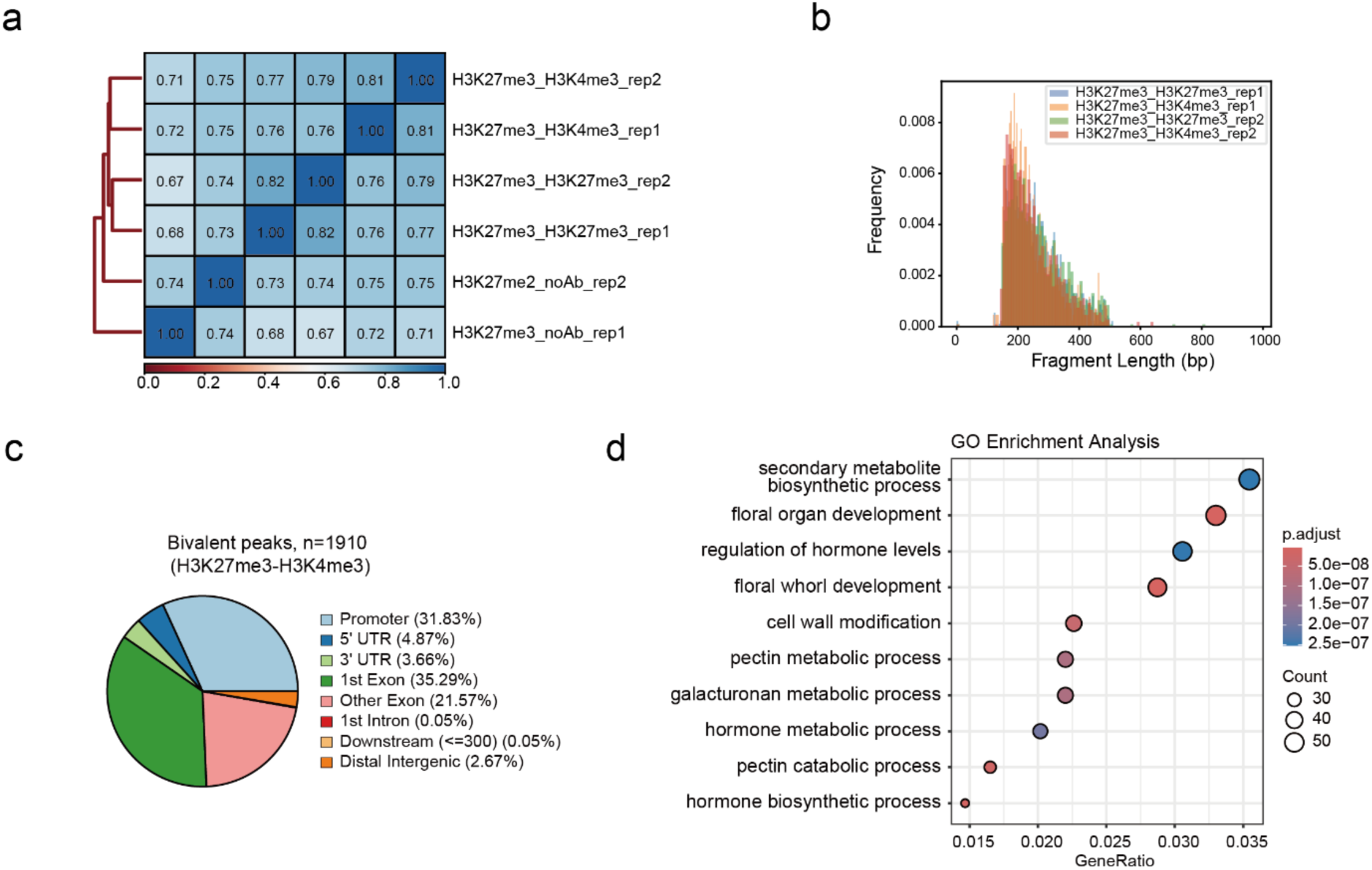
Quality control of sequential ChIP-seq. **a.** Pairwise Spearman correlation heatmap showing the similarity among sequential ChIP-seq samples from 14-day-old Col-0 seedlings, including H3K27me3-H3K27me3, H3K27me3-H3K4me3, and H3K27me3-noAb (no antibody) control samples, with two biological replicates for each condition. Colours represent Spearman correlation coefficients. **b.** Fragment-length distributions for H3K27me3-H3K27me3 and H3K27me3-H3K4me3 sequential ChIP-seq libraries across biological replicates. **c.** Genomic annotation of H3K27me3-H3K4me3 bivalent peaks identified by sequential ChIP-seq (n=1910). Pie chart shows the proportion of peaks assigned to promoters, UTRs, exons, introns, downstream regions, and distal intergenic regions. **d.** Gene Ontology (GO) Biological Process enrichment analysis of genes associated with H3K27me3-H3K4me3 bivalent peaks (n=1800). Dot size represents gene count, and colour indicates the adjusted *P* value.

**S Fig. 11.**
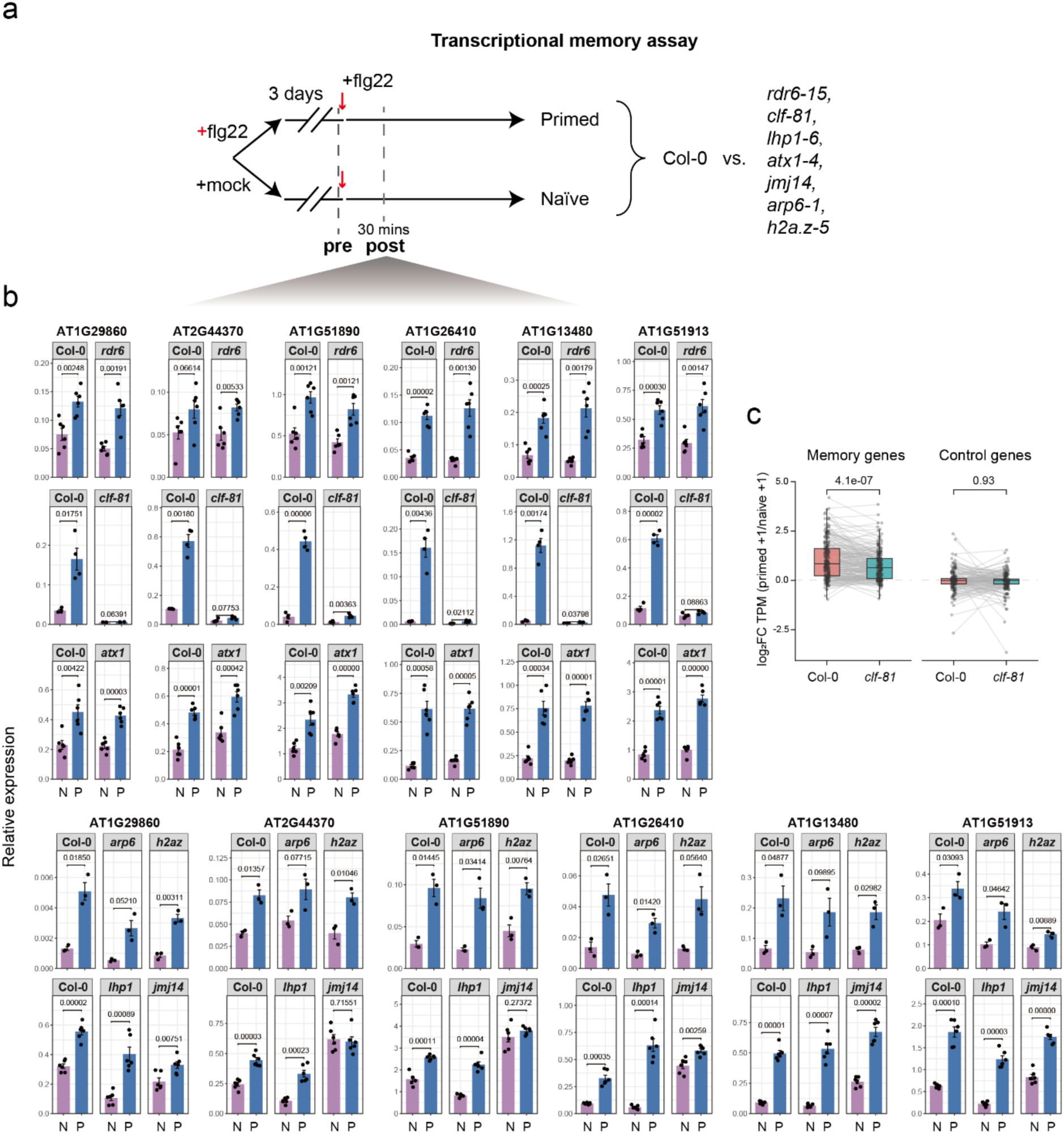
Transcriptional memory analysis of distinct chromatin mutants. **a.** Schematic of the transcriptional memory assay used to compare Col-0 with the indicated mutant backgrounds. Seedlings were primed with flg22 (primed) or mock (naïve), allowed to recover for 3 days, and then challenged with flg22. Samples were collected immediately before the recurrent flg22 treatment (pre) and 30 mins after treatment (post). **b.** Bar plots showing IPP2-normalized relative expression of six representative memory genes in Col-0 and the indicated chromatin mutant backgrounds under naïve (N) and primed (P) conditions. Gene expression was measured 30 mins after recurrent flg22 treatment. Bars represent mean relative expression ± SEM, and individual points represent biological replicates (n = 3, 4 or 6). Statistical significance between naïve and primed samples within each genotype was assessed using two-sample t-tests, with *P* values indicated above the corresponding comparisons. **c.** Paired box plots showing the priming response, calculated as log_2_[(primed TPM + 1)/(naïve TPM + 1)], for memory genes and control genes in Col-0 and *clf-81*. Grey lines connect the same genes between genotypes. Box plots show the median and interquartile range, with whiskers extending to 1.5x the interquartile range. *P* values were calculated using a paired Wilcoxon signed-rank test.

**S Fig. 12.**
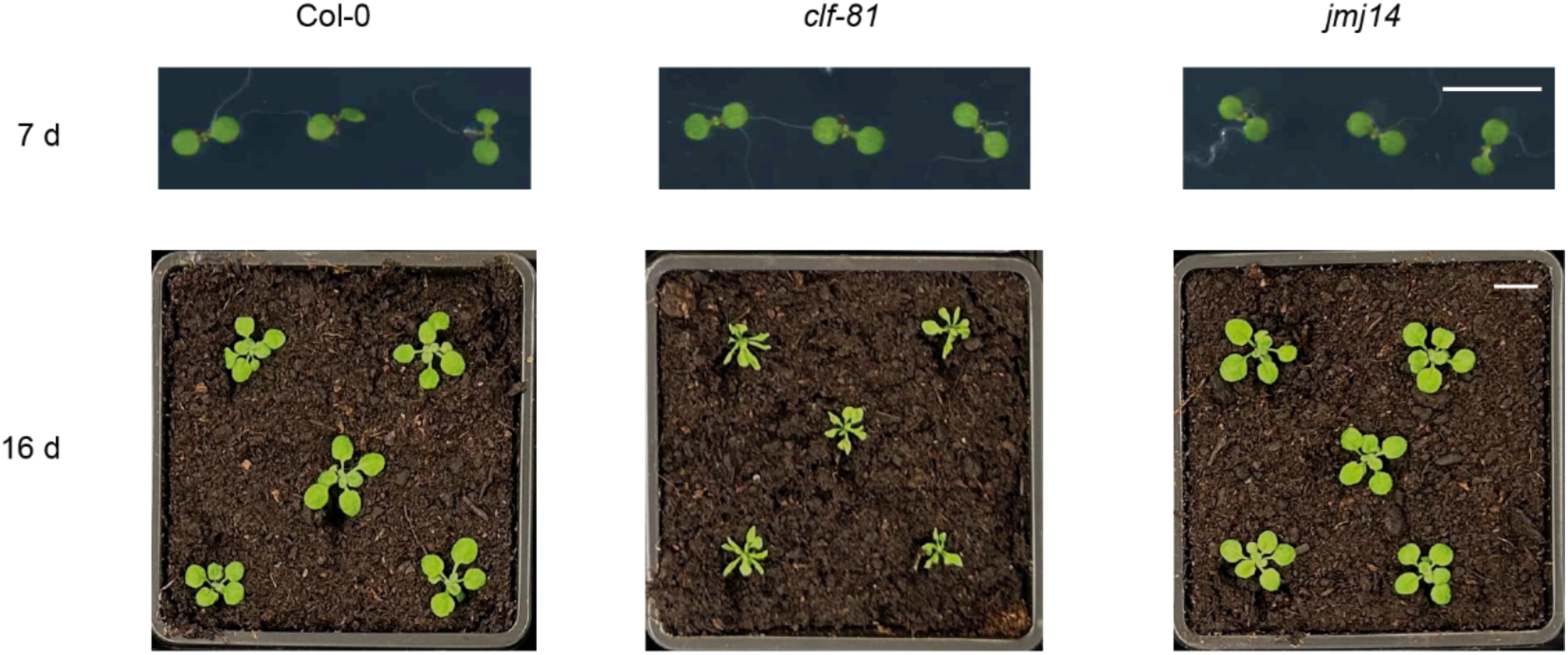
Developmental phenotypes of Col-0, *clf-81*, and *jmj14*. Representative images of Col-0, *clf-81*, and *jmj14* plants at 7 and 16 days after germination, showing developmental phenotypes at early seedling and later vegetative stages. Scale bars, 1 cm.

**S Fig. 13.**
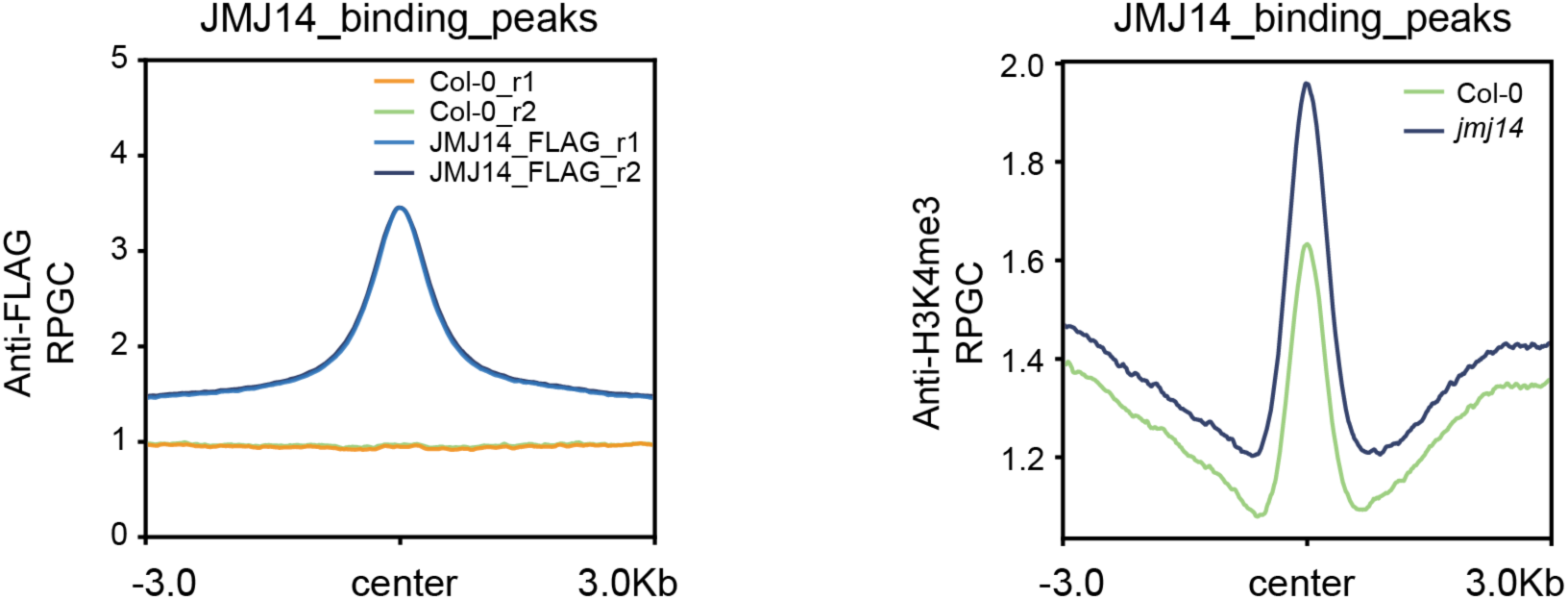
Expected profiling of published ChIP-seq datasets (GSE204681). Metaplots showing anti-FLAG ChIP-seq signal over JMJ14-binding peaks, with enrichment in JMJ14-FLAG samples relative to Col-0 controls across two biological replicates (left), and H3K4me3 ChIP-seq signal over JMJ14-binding peaks in Col-0 and *jmj14* (right).

**S Fig. 14.**
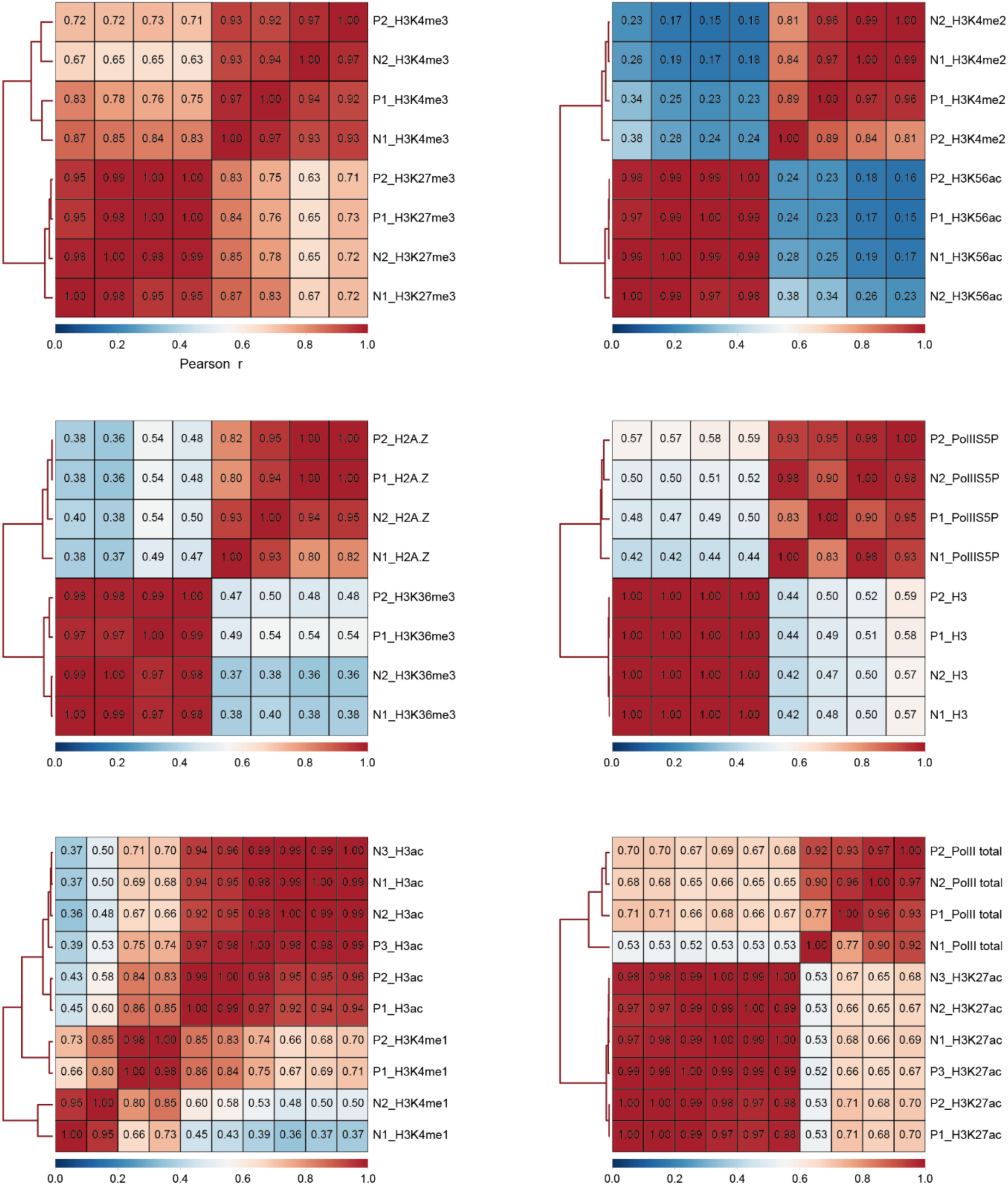
Correlation between biological replicates in ChIP-seq data. Pairwise Pearson correlation heatmaps showing the reproducibility of ChIP-seq datasets for the indicated chromatin marks and Pol II profiles in naïve (N) and primed (P) samples collected at the end of the 3-day recovery period following priming treatment. N1, N2, and N3 represent biological replicates of the naïve condition, whereas P1, P2, and P3 represent biological replicates of the primed condition. Colours and values represent Pearson correlation coefficients (r), with hierarchical clustering illustrating similarity among samples.

**S Fig. 15.**
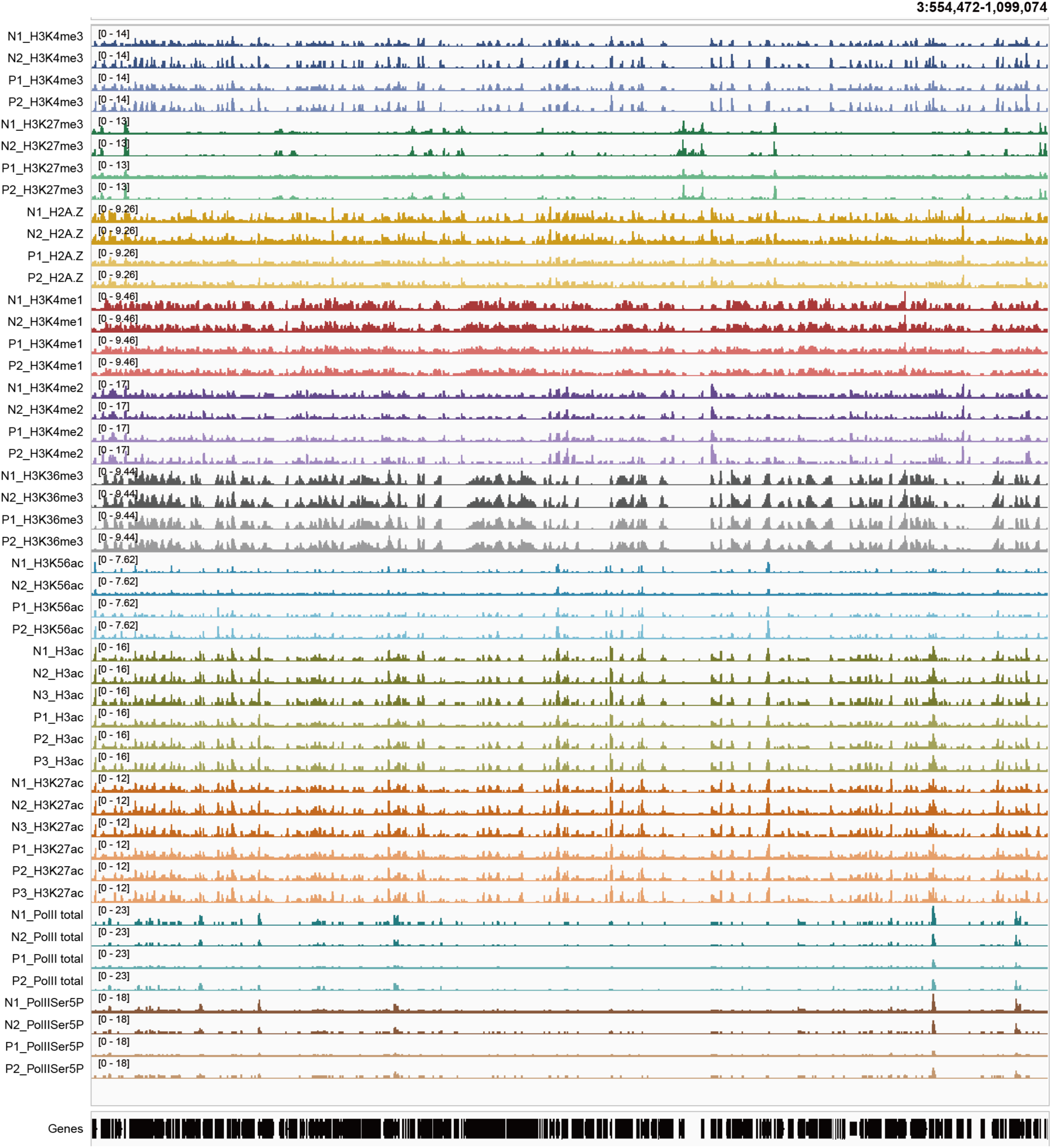
IGV genome browser screenshots showing all ChIP-seq biological replicates from naïve and primed samples across a randomly selected genomic region.

**S Fig. 16.**
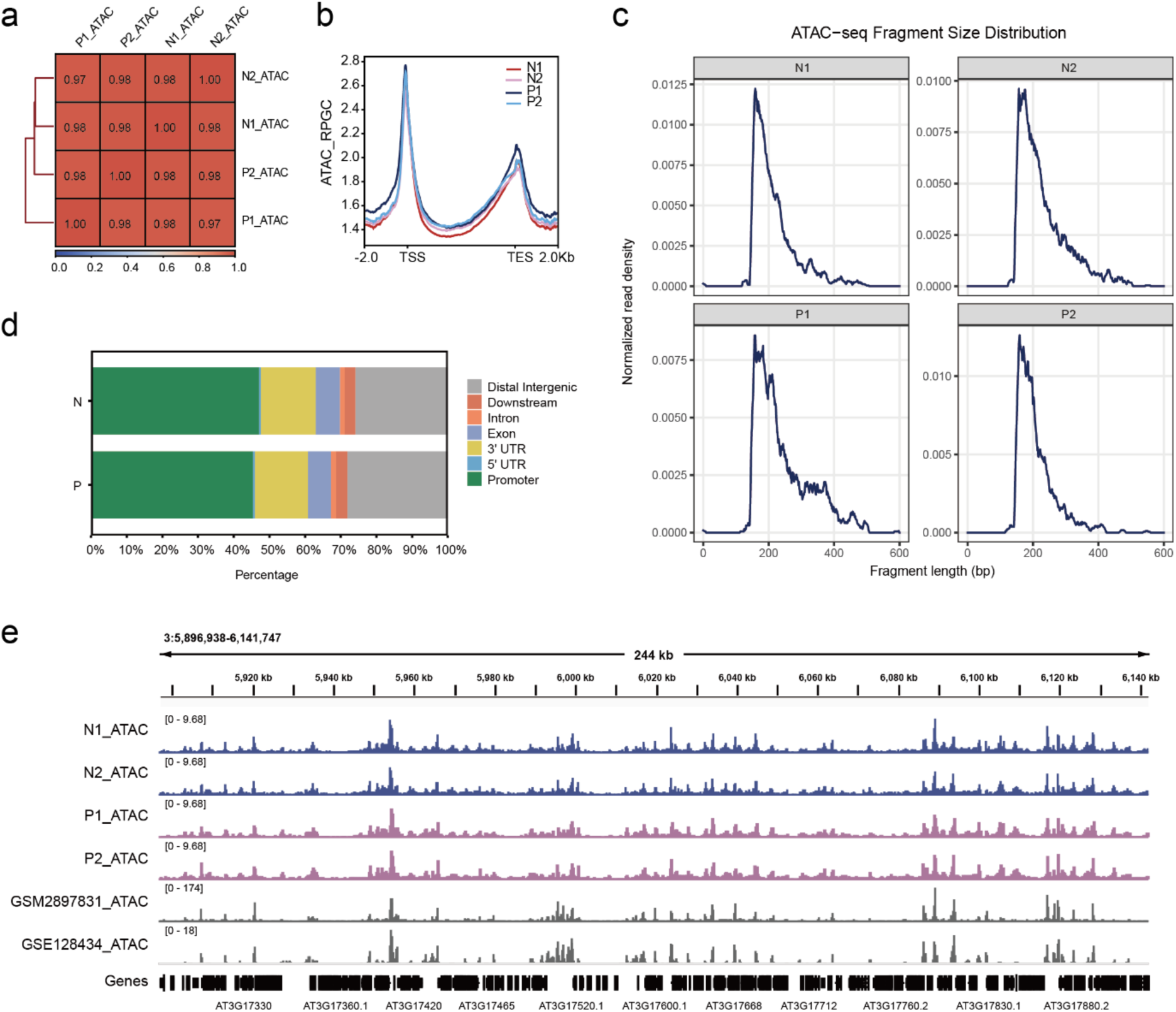
Quality control of ATAC-seq data. **a.** Spearman correlation analysis of ATAC-seq data across biological replicates. Two biological replicates were analyzed for each condition (N1 and N2, naïve; P1 and P2, primed). Color intensity indicates Spearman correlation coefficients. **b.** Metaplots showing ATAC-seq signals across all genes for biological replicates under naïve (N) and primed (P) conditions. **c.** ATAC-seq fragment size distributions for naïve (N1, N2) and primed (P1, P2) samples. Curves show normalized read density as a function of fragment length.**d.** Genomic annotation of ATAC-seq peaks in naïve (N) and primed (P) samples, with biological replicates merged separately within each condition. Peaks were assigned to genomic features (promoter, UTRs, exons, introns, downstream, and distal intergenic regions), and the percentage of peaks in each category is shown. **e.** IGV genome browser screenshots showing all ATAC-seq biological replicates from this study together with two published ATAC-seq datasets (GSM2897831 and GSE128434) across a randomly selected genomic region.

**S Fig. 17.**
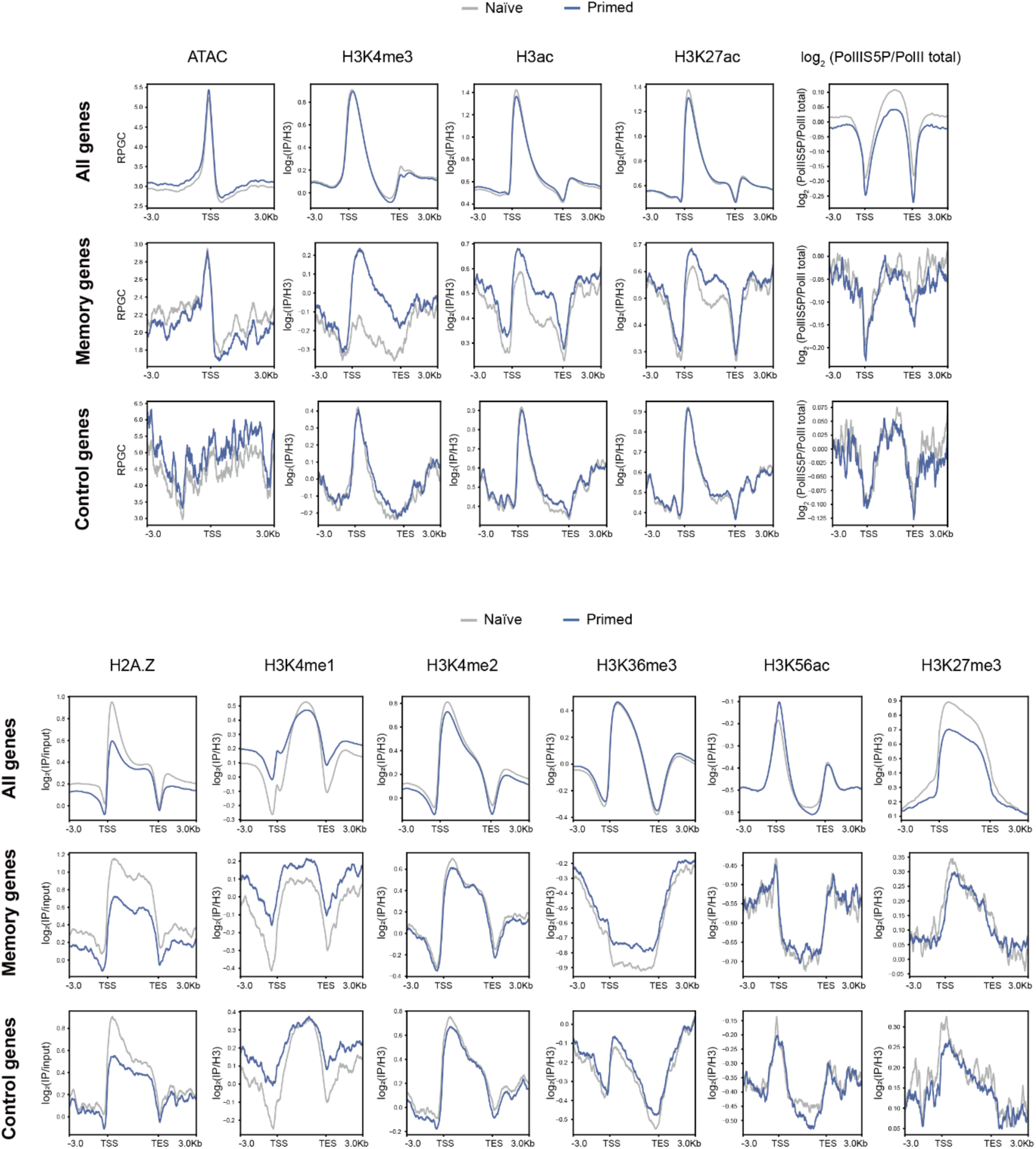
Chromatin profiles of all genes, memory genes, and control genes in naïve and primed conditions. Metaplots showing ATAC-seq and ChIP-seq profiles across all genes (top), memory genes (middle), and one set of transcriptionally equivalent non-memory control genes (bottom) under naïve (grey) and primed (blue) conditions after 3 days of recovery and before recurrent flg22 treatment. H3K4me3, H3ac, H3K27ac, H3K4me1, H3K4me2, H3K36me3, H3K56ac, and H3K27me3 signals were normalized to H3; H2A.Z was normalized to input; Pol II Ser5P was normalized to total Pol II; and ATAC-seq signal is shown as RPGC-normalized accessibility. Profiles were generated from data merged from two or three biological replicates per condition.

**S Fig. 18.**
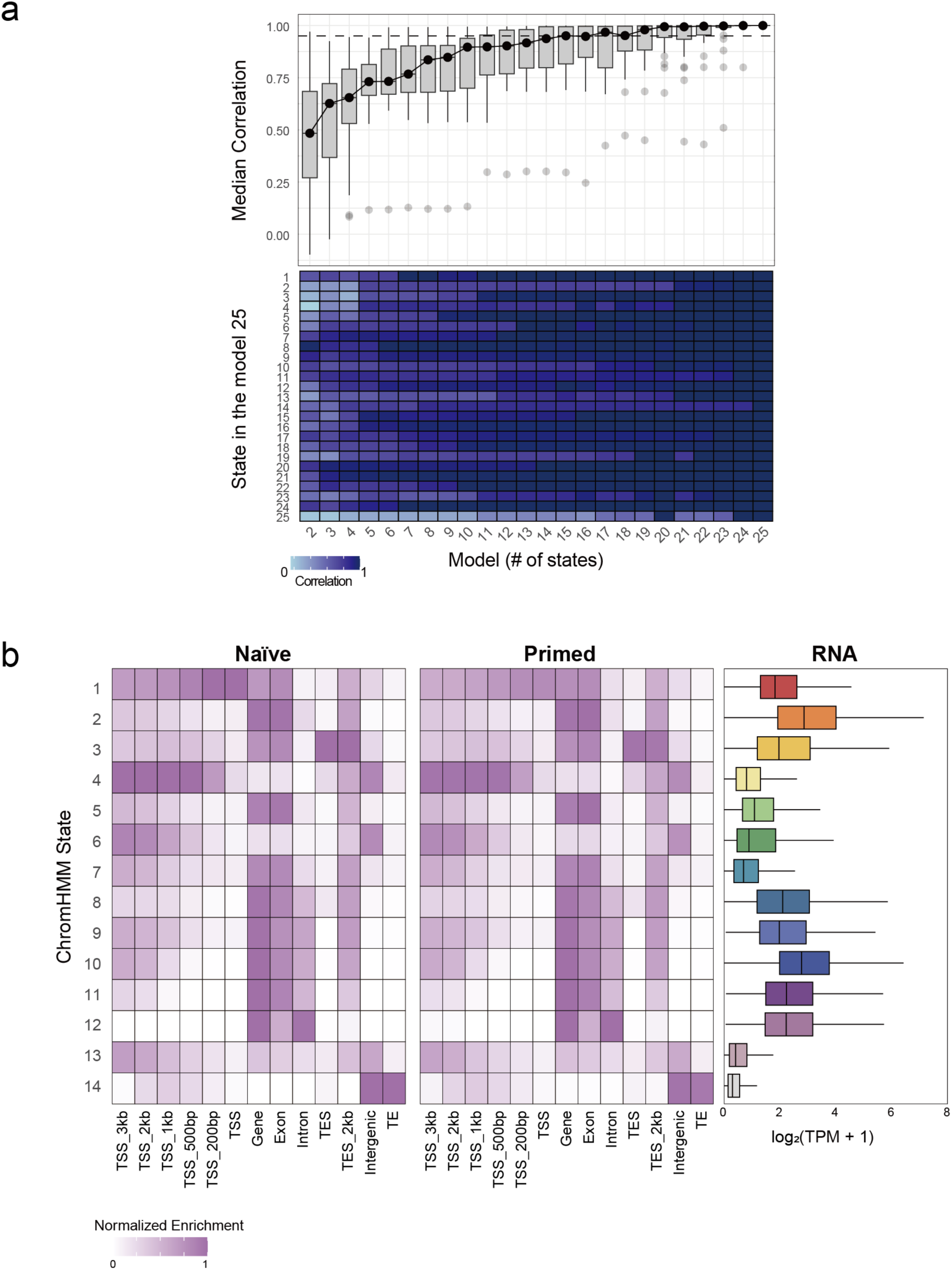
ChromHMM model selection and chromatin-state annotation. **a.** ChromHMM model comparison across models containing 2-25 chromatin states. Top, distribution of state correlations for each model relative to the 25-state model, with black points indicating the median correlation and the dashed horizontal line indicating a correlation threshold of 0.95. Bottom, heatmap showing the correspondence between individual states in the 25-state model and states in models containing fewer states. **b.** Genomic annotation of the 14-state ChromHMM model in naïve and primed conditions. Heatmaps show normalized enrichment of each chromatin state across genomic features, including regions surrounding the transcription start site (TSS), gene bodies, exons, introns, transcription end sites (TES), intergenic regions, and transposable elements (TEs). Right, RNA expression distributions for genes associated with each chromatin state, shown as log_2_(TPM + 1).

**S Fig. 19.**
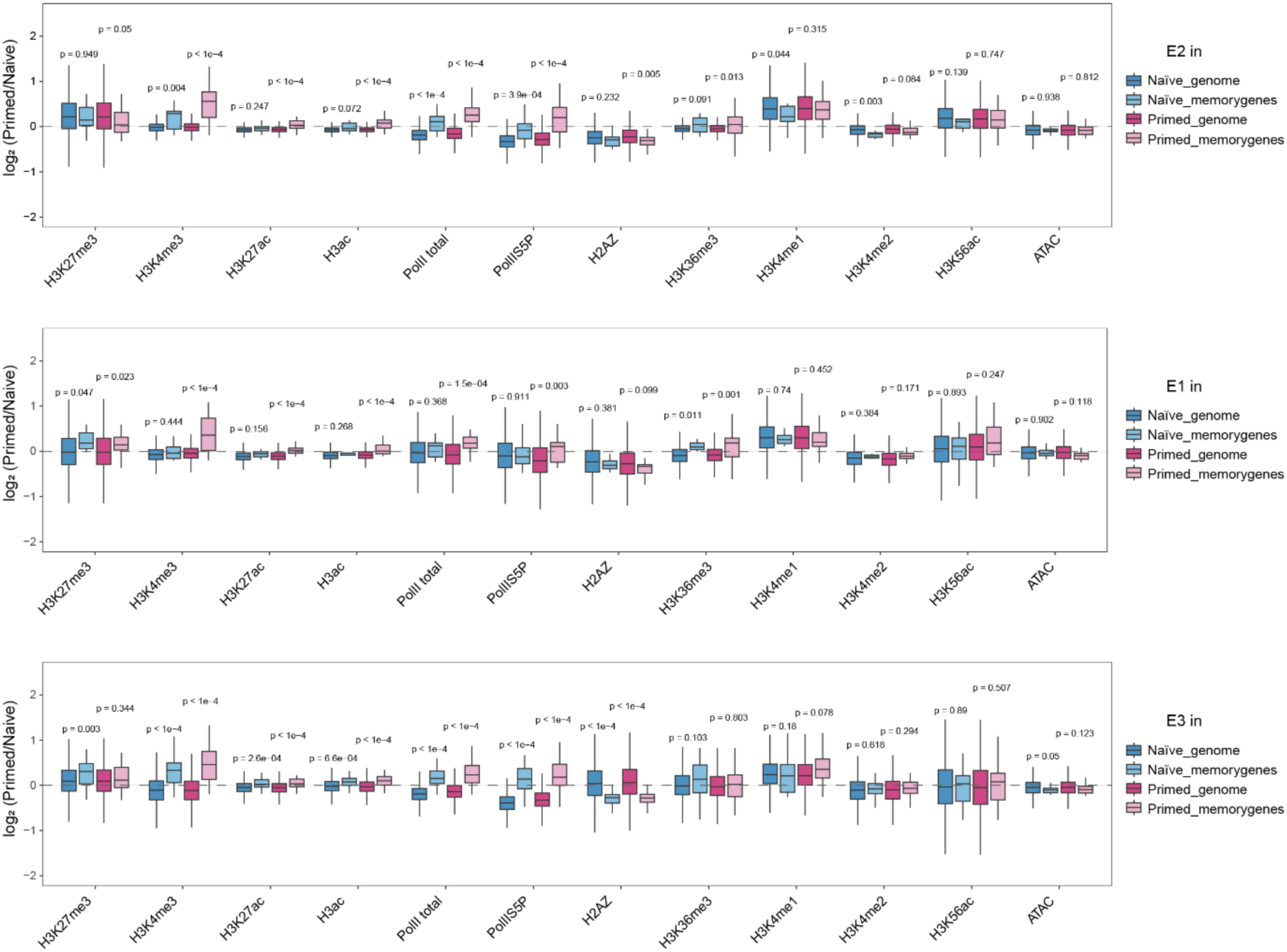
Chromatin feature changes within selected ChromHMM states in naïve and primed conditions. Boxplots showing log_2_ fold-change values (primed/naïve) for the indicated chromatin features and ATAC-seq accessibility within ChromHMM states E2 (top), E1 (middle), and E3 (bottom), comparing genome-wide intervals with intervals associated with memory genes. Features include H3K27me3, H3K4me3, H3K27ac, H3ac, total Pol II, Pol II Ser5P, H2A.Z, H3K36me3, H3K4me1, H3K4me2, H3K56ac, and ATAC-seq. The top, centre line, and bottom of each box represent the upper quartile, median, and lower quartile, respectively. *P* values were calculated using Wilcoxon rank-sum tests and are indicated above the corresponding comparisons.

**S Fig. 20.**
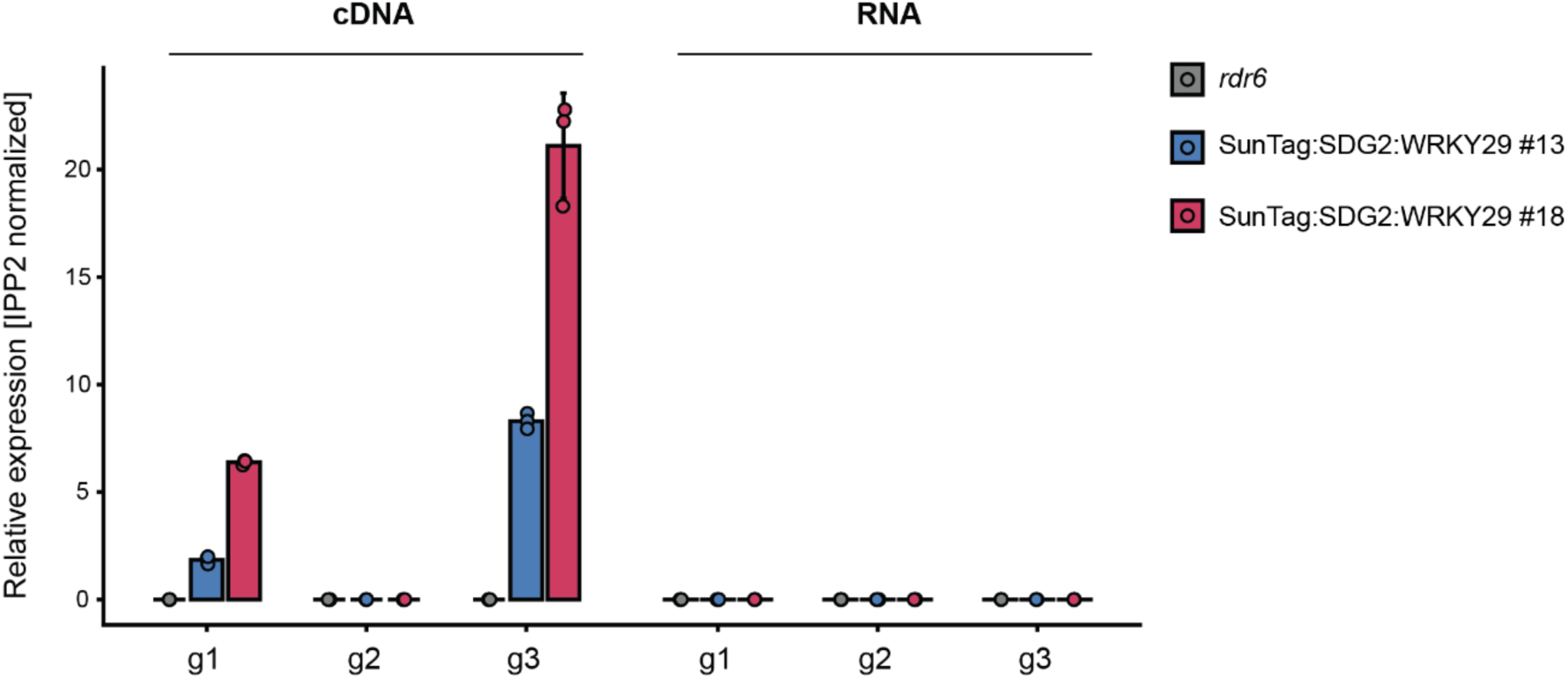
Validation of guide RNA expression in SunTag:SDG2:WRKY29 lines. Bar plots showing IPP2-normalized expression of *WRKY29* guide RNAs g1, g2, and g3 in *rdr6* and two independent SunTag:SDG2:WRKY29 lines (#13 and #18). Expression was measured from scaffold-specific cDNA (left), with corresponding RNA samples included as negative controls (right) to confirm that the detected signal was derived from reverse-transcribed RNA rather than contaminating DNA. Bars represent mean expression, with individual replicates shown as points.

**S Fig. 21.**
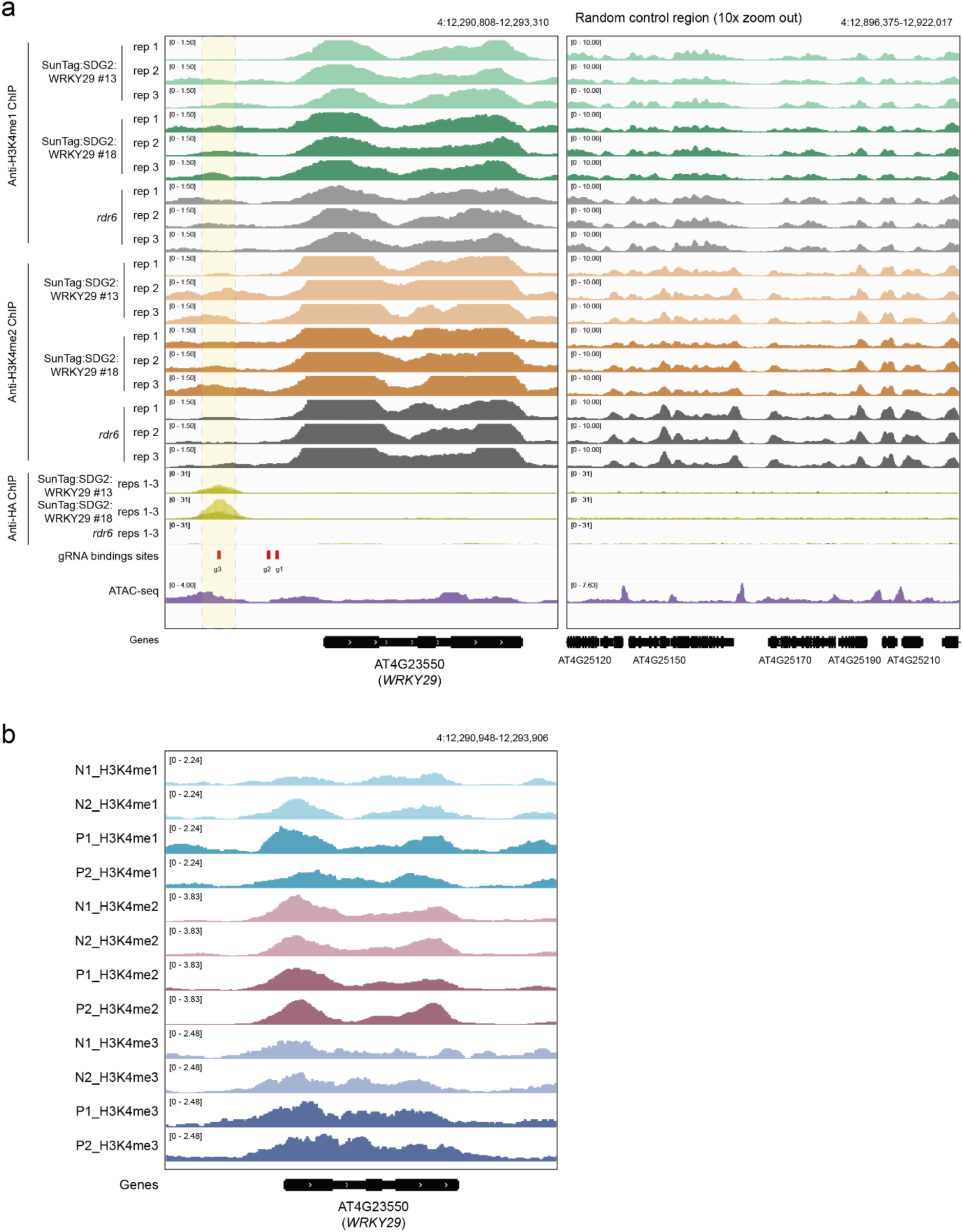
H3K4me enrichment at targeted *WRKY29* and their dynamics upon priming. **a.** IGV genome browser snapshots showing H3K4me1/2 enrichment at the *WRKY29* locus in two independent SunTag:SDG2:WRKY29 lines (#13 and #18) and the *rdr6* control, together with anti-HA ChIP-seq signal marking SunTag occupancy, gRNA-binding sites, and ATAC-seq accessibility. Three biological replicates are shown for H3K4me1/2 ChIP-seq, while the anti-HA ChIP-seq signal represents the overlay of three biological replicates. The yellow shaded region indicates the targeted gRNA-binding region. A random control region is shown on the right. **b.** IGV genome browser snapshots showing H3K4me1, H3K4me2 and H3K4me3 profiles at the *WRKY29* locus in individual biological replicates under naïve and primed conditions. N1 and N2 indicate naïve replicates, P1 and P2 indicate primed replicates.

**S Fig. 22.**
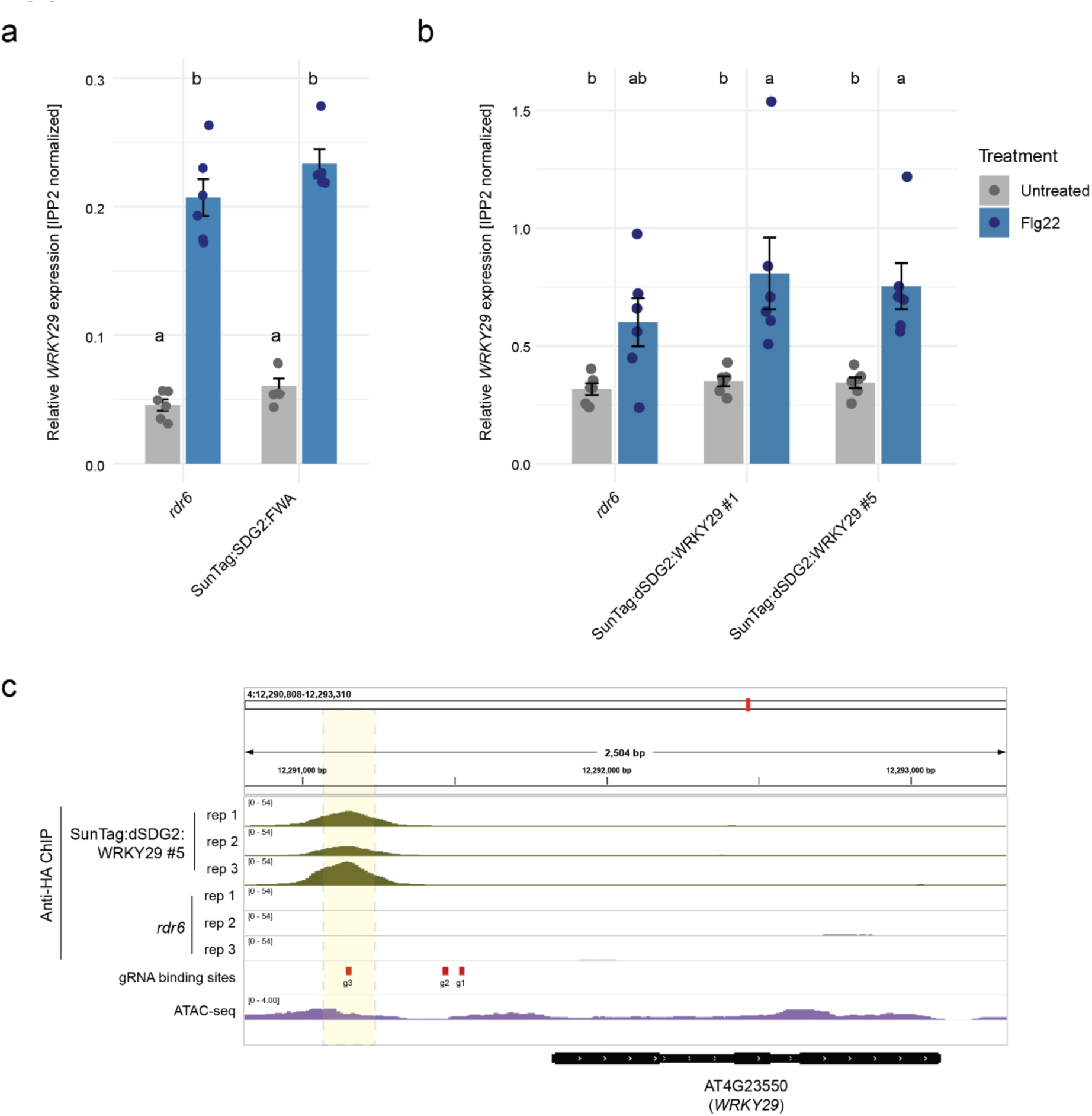
Negative controls for SunTag-SDG2 or dSDG2-mediated epigenetic editing at *WRKY29*. **a.** Bar plots showing IPP2-normalized *WRKY29* expression in *rdr6* and the SunTag:SDG2:FWA control line under untreated and flg22-treated conditions (n = 6 biological replicates). Bars represent mean expression, with individual biological replicates shown as points. Different letters indicate statistically significant differences among groups, determined by one-way ANOVA followed by Tukey’s HSD test (*P* < 0.05). **b.** Bar plots showing IPP2-normalized *WRKY29* expression in *rdr6* and two independent SunTag:dSDG2:WRKY29 lines (#1 and #5) under untreated and flg22-treated conditions (n = 6 biological replicates). Bars represent mean expression, with individual biological replicates shown as points. Different letters indicate statistically significant differences among groups, determined by one-way ANOVA followed by Tukey’s HSD test (*P* < 0.05). **c.** IGV genome browser view showing anti-HA ChIP-seq signal at the *WRKY29* locus in SunTag:dSDG2:WRKY29 line #5 and *rdr6* controls across three biological replicates. gRNA-binding sites and ATAC-seq accessibility are shown. The yellow shaded region indicates the targeted gRNA-binding region.

**S Fig. 23.**
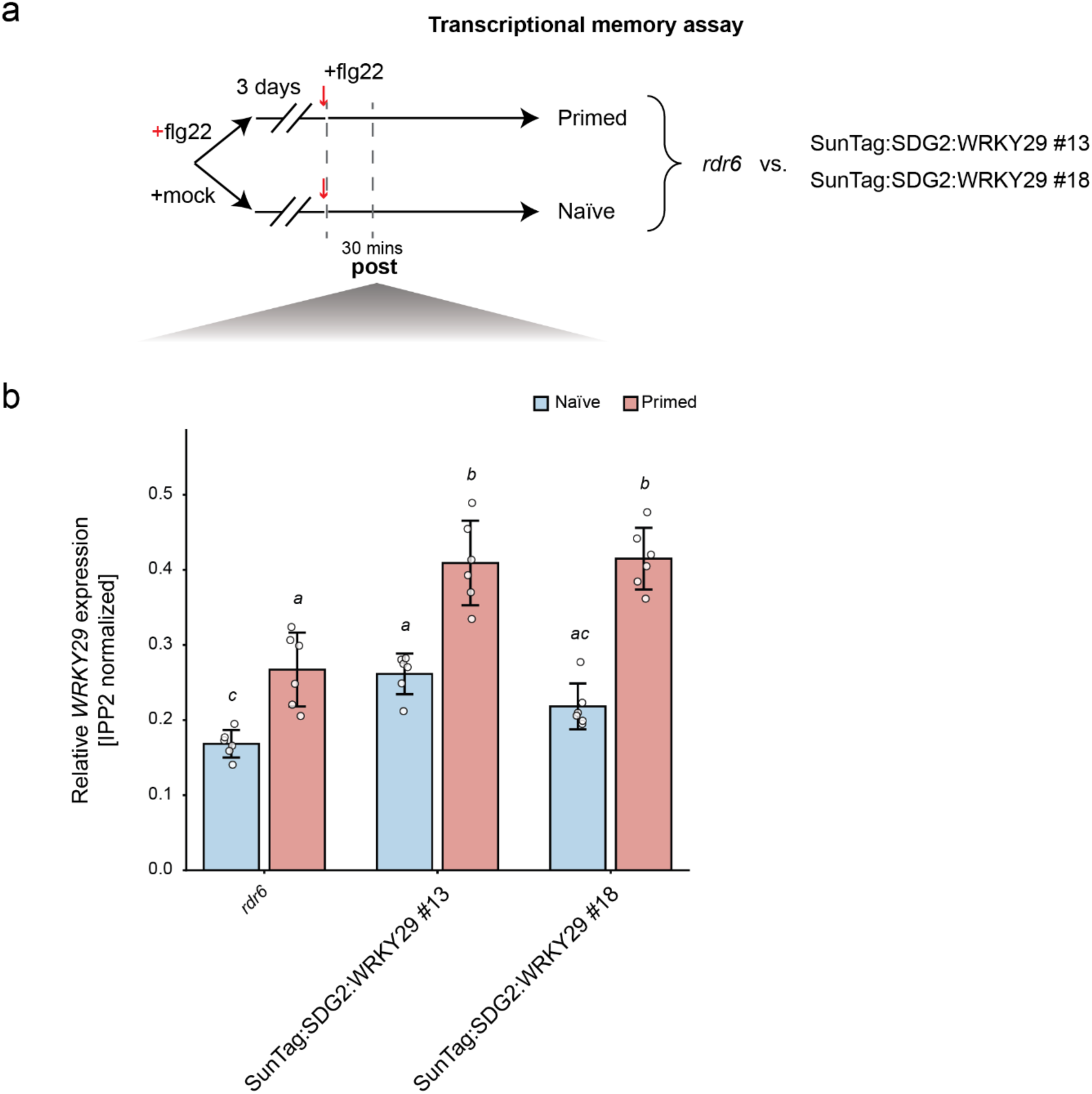
Transcriptional memory in SunTag:SDG2:WRKY29 lines. **a.** Schematic of the transcriptional memory assay used to compare *rdr6* with two independent SunTag:SDG2:WRKY29 lines (#13 and #18). Seedlings were primed with flg22 (primed) or mock (naïve), allowed to recover for 3 days, and then challenged with flg22. Samples were collected 30 mins after the recurrent flg22 treatment (post), with six biological replicates per genotype and condition. **b.** Bar plots showing IPP2-normalized *WRKY29* expression in *rdr6* and two independent SunTag:SDG2:WRKY29 lines (#13 and #18) under naïve (N) and primed (P) conditions, measured 30 mins after recurrent flg22 treatment. Bars represent mean expression ± SEM, with individual biological replicates shown as points. Different letters indicate statistically significant differences among groups, determined by one-way ANOVA followed by Tukey’s HSD test (*P* < 0.05).

**S Fig. 24.**
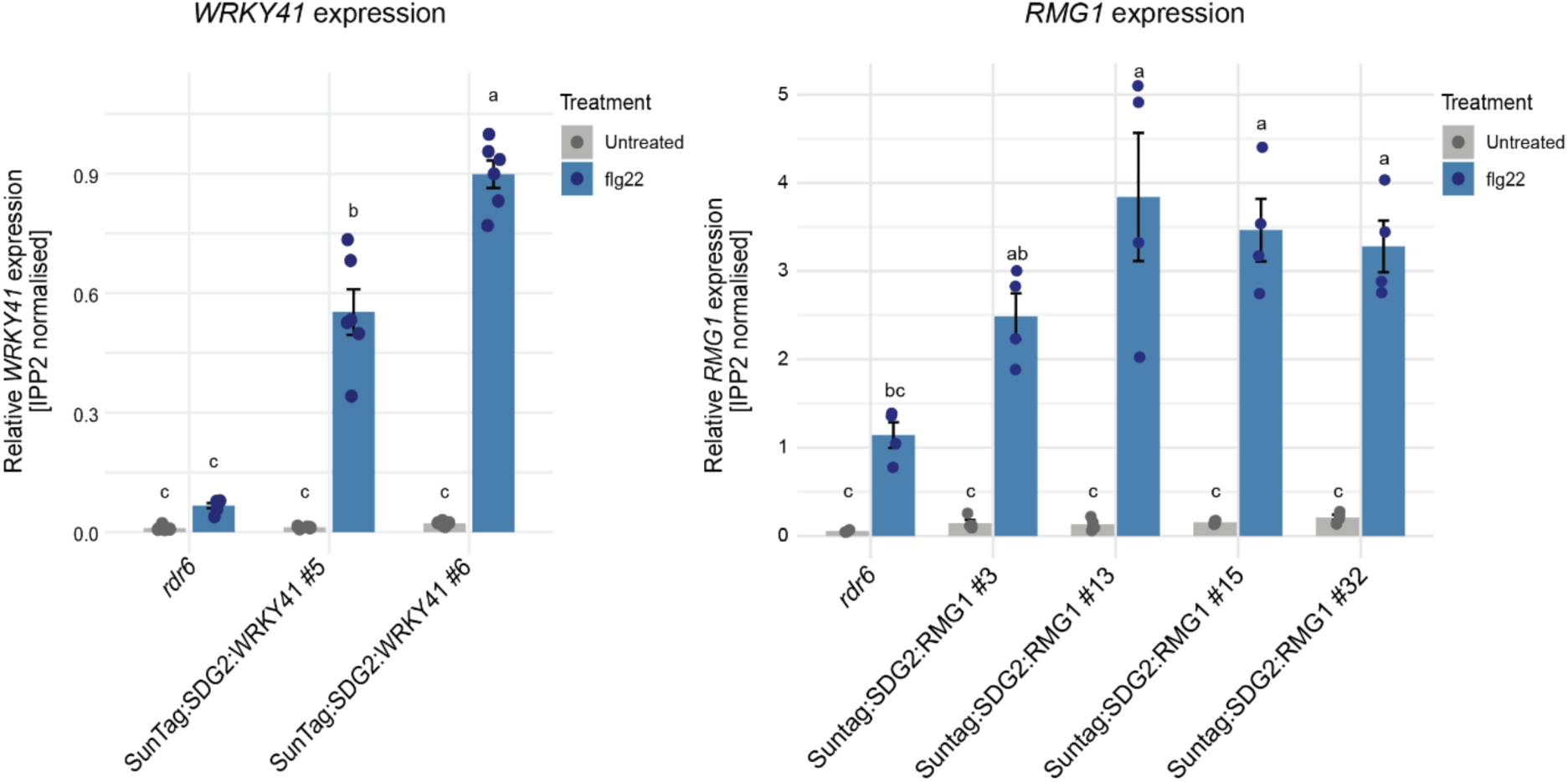
Validation of SunTag:SDG2 targeting at *WRKY41* and *RMG1*. Bar plots showing IPP2-normalized expression of *WRKY41* (left) and *RMG1* (right) in *rdr6* and independent SunTag:SDG2 lines targeting the indicated loci under untreated and flg22-treated conditions. For *WRKY41*, n = 6 biological replicates per genotype and treatment; for *RMG1*, n = 4 biological replicates per genotype and treatment. Bars represent mean expression with individual biological replicates shown as points. Different letters indicate statistically significant differences among groups, determined by one-way ANOVA followed by Tukey’s HSD test (*P* < 0.05).

